# Self-contrastive learning enables interference-resilient and generalizable fluorescence microscopy signal detection without interference modeling

**DOI:** 10.1101/2025.04.08.645087

**Authors:** Fengdi Zhang, Ruqi Huang, Meiqian Xin, Haoran Meng, Danheng Gao, Ying Fu, Juntao Gao, Xiangyang Ji

**Affiliations:** Shenzhen International Graduate School, Tsinghua University, Shenzhen, 518055, China; Pengcheng Laboratory, Shenzhen, 518055, China; Center for Synthetic and Systems Biology, Tsinghua University, Beijing, 100084, China; Institute of TCM-X, Tsinghua University, Beijing, 100084, China; Changchun Institute of Optics, Fine Mechanics and Physics, Chinese Academy of Sciences, Changchun, 130033, China; School of Computer Science and Technology, Beijing Institute of Technology, Beijing, 100081, China; Department of Automation, Tsinghua University, Beijing, 100084, China

## Abstract

Every weak signal in fluorescence microscopy may contain critical biological information. However, the interference resilience required to detect such signals has traditionally relied on task-specific interference modeling, which limits generalizability. Here, we present a self-contrastive learning-based signal detection solution that achieves interference resilience without the need for interference modeling, thereby offering high generalizability. The method, DEep PAttern Fitting (DEPAF), is a module that contrasts asynchronously generated data views from asymmetric model paths to extract signals from interference, while incorporating highly parallel signal recognition and localization in the process. Benchmarking shows that DEPAF substantially improves signal detection performance across diverse imaging modalities and dimensions, especially under challenging conditions such as low SNRs and ultra-high signal densities. It is also compatible with, and consistently enhances the performance of various imaging techniques, such as super-resolution imaging, spatial transcriptomic imaging, and two-photon calcium imaging. Notably, DEPAF relies only on image patches with signal patterns and one tunable hyperparameter to adapt to new tasks, making it accessible to users without domain-specific expertise and lowering the barrier to broader adoption. DEPAF is expected to advance the signal-centric fluorescence microscopy techniques and inspire further advancements, especially in the era of image-based multi-omics.

## Introduction

In the world of fluorescence microscopy, the interplay between light and molecular signals at the microscopic scale reveals a detailed and dynamic picture of cellular and molecular activities, playing a crucial role in analyzing and understanding biological dynamics and molecular mechanisms. Signal detection—determining the presence of signals and localizing them—is a fundamental step in many advanced fluorescence microscopy techniques. Examples abound, such as localization-based super-resolution imaging [1], omics analysis [2], and neural activity analysis [3]. However, with the growing demand for higher resolution, greater throughput, and broader applicability, signal detection is increasingly challenged by various forms of interference, including decreased signal-to-noise ratio (SNR) due to reduced photon budgets [4–6], strong fluorescence background caused by optical scattering and non-specific staining [7, 8], and severe signal overlap resulting from increased labeling density [4, 9].

Although adjustments in experimental design can partially alleviate the above issues, such as increasing laser excitation intensity, introducing autofluorescence quenching steps, or sparsifying the labels in space and time, these approaches are essentially trade-offs. They often come with side effects, including causing phototoxicity and damaging live cells [4–6, 10], degrading sample quality [7, 8], and compromising the spatiotemporal resolution of observations [2, 4]. Therefore, solutions in data processing are crucial. Detection methods that are resilient to interference and generalizable across various types of fluorescence microscopy signals stand at the fulcrum of this field.

However, challenges persist, as interference-resilience typically depends on interference modeling for specific imaging modalities. This dependency undermines the generalizability of detection methods, leading to the lack of a method that is both generalizable and interference-resilient (Fig. 1a). For instance, self-supervised denoising methods [6, 10–14] enhance signals by leveraging the unpredictability of noise and redundancy of signals. These methods achieve image denoising without modeling the noise but cannot distinguish background interference or detect signals. Conventional statistical methods [15–26] estimate signals from imaging data by fitting explicit probabilistic models without needing interference modeling. Unfortunately, their performance worsens under severe signal overlap, strong noise, or complex backgrounds, showing poor robustness against interference. Supervised learning methods [27–29] rely heavily on labeled data, limiting their generalizability to tasks requiring high signal density or extremely high accuracy, where labeled data is often unavailable [1, 22]. Simulation learning methods [4, 9, 30–33] use detailed interference models to simulate labeled data for supervised learning, demonstrating high interference resistance. Yet, these methods must be specifically tailored to each imaging modality, limiting their generalizability. Table 1 summarizes the characteristics of these methods. Therefore, developing an accessible fluorescence microscopy signal detection method that is both generalizable and resistant to interference remains a challenge, both conceptually and practically, for multi-modal biological signal analysis and understanding broader biological phenomena.

**Table 1.** Comparison of fluorescence microscopy signal processing methods. (A) Signal detection, (B) Interference-resilient, (C) No external ground truth needed, (D) No interference modeling needed.

| Method | A | B | C | D |
| --- | --- | --- | --- | --- |
| Self-supervised denoising [6, 10–14] | ✗ | ✓ | ✓ | ✓ |
| Conventional statistical [15–26] | ✓ | ✗ | ✓ | ✓ |
| Supervised learning [27–29] | ✓ | ✓ | ✗ | ✓ |
| Simulation learning [4, 9, 30–33] | ✓ | ✓ | ✓ | ✗ |
| **Self-contrastive learning (DEPAF)** | ✓ | ✓ | ✓ | ✓ |

**Fig. 1.**
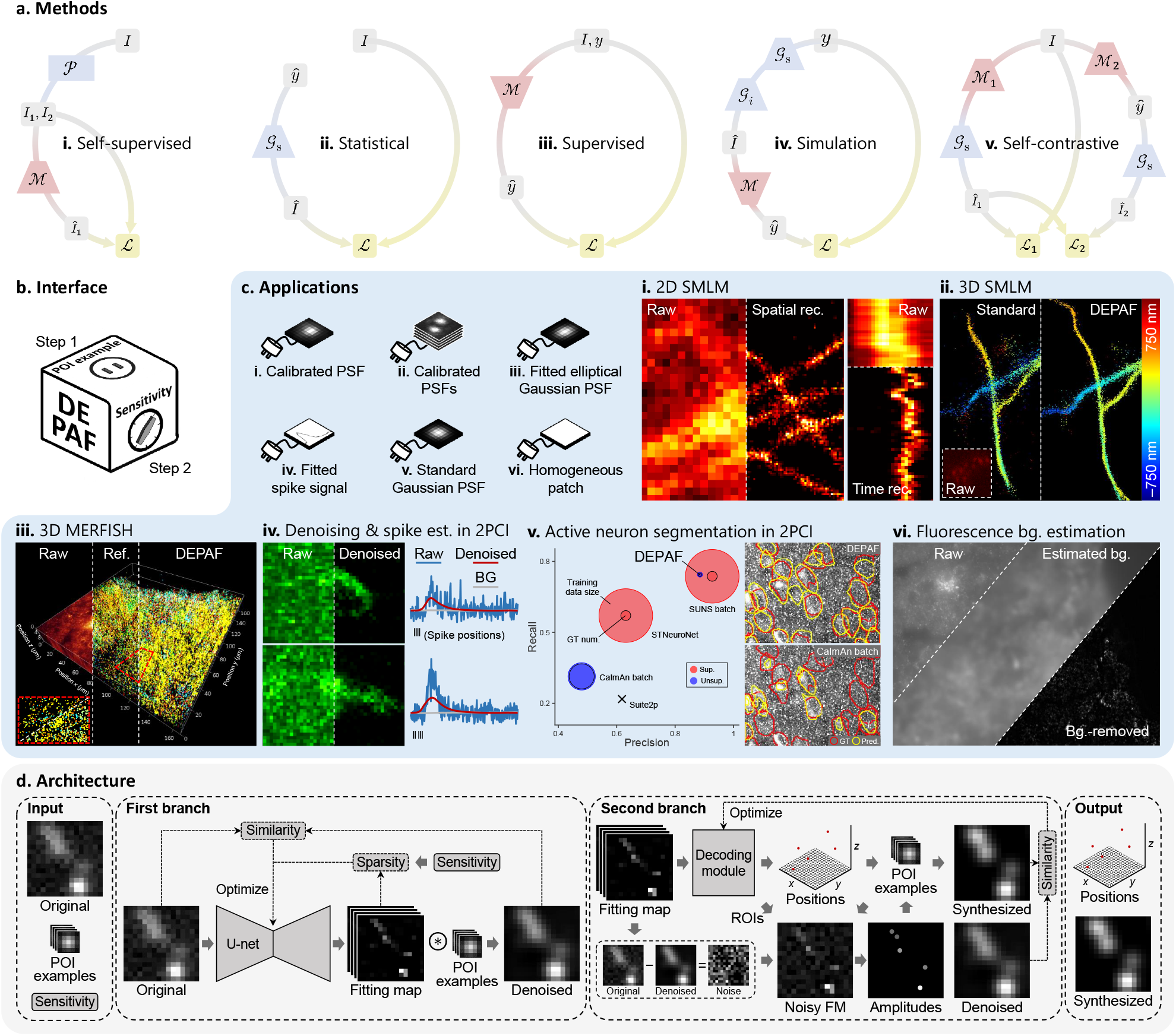
General overview of DEPAF. **a**, Fluorescence microscopy signal processing methods. **i**. Self-supervised denoising methods. Noise-independent but signal-consistent image pairs *I*_1_, *I*_2_ are derived from input images *I* via pretext tasks 𝒫. *I*_1_ are fed into denoising model ℳ to predict clean images *Î*_1_. ℳ is optimized by loss ℒ between *Î*_1_ and *I*_2_. **ii**. Conventional statistical methods. Detection results *ŷ* are passed through signal model 𝒢_*s*_ to generate synthesized data *Î*. *ŷ* are optimized by loss ℒ between *Î* and input images *I*. **iii**. Supervised learning methods. Input images *I* are processed by model ℳ to produce detection results *ŷ*. ℳ is optimized by loss ℒ between *ŷ* and external ground truth *y*. **iv**. Simulation learning methods. Starting from generated ground truth *y*, ideal signals are generated via 𝒢 _*s*_ and corrupted using interference model 𝒢 _*i*_ to obtain simulated images *Î*. *Î* are input to model ℳ to get detection results *ŷ*. ℳ is optimized by loss ℒ between *y* and *ŷ*. **v**. Self-contrastive learning methods *applied in this work*. Based on input images *I*, detection results *ŷ* and two views, *Î*_1_ and *Î*_2_, each containing a mixture of signals with noisy signals and signals with noise, respectively, are obtained through asymmetric and asynchronous model paths, involving models ℳ_1_, ℳ_2_, and signal model 𝒢_*s*_ *without* requiring any interference models. ℳ_1_ is first optimized by loss ℒ _1_ between *Î*_1_ and *I*, followed by optimizing ℳ_2_ using loss ℒ _2_ between *Î*_1_ and *Î*_2_. **b**, DEPAF provides a streamlined interface, enabling users to achieve diverse fluorescence microscopy signal detection by simply replacing the POI examples (step 1) and adjusting a single hyperparameter corresponding to detection sensitivity (step 2). **c**. The POI examples used for different applications, along with the corresponding result diagrams. **i**. DEPAF elevates the temporal resolution of 2D SMLM to the millisecond level without compromising the expected spatial resolution. Rec., reconstruction. **ii**. In 3D SMLM, DEPAF maintains stable and high-quality reconstruction performance even under low photon counts and high blinking densities. **iii**. In 3D MERFISH imaging data with strong background, DEPAF notably improves RNA spot signal detection rates and quantitative accuracy. Ref., reference. **iv**. For low-SNR two-photon calcium imaging of neurons, DEPAF achieves efficient denoising, background separation, and detection of weak neural activity. Est., estimation; 2PCI, two-photon calcium imaging; BG, background. **v**. Leveraging the denoised results from **iv**, DEPAF, as an unsupervised source extraction-based method, enables high-performance active neuron segmentation with minimal data requirements, clearly outperforming other such methods, and is even competitive with top-performing supervised methods. GT num., required number of ground-truth labels; Sup., supervised; Unsup., unsupervised; Pred., prediction. **vi**. DEPAF can estimate arbitrary inhomogeneous fluorescence backgrounds, adapting to varying SNR levels and background variation scales. Bg., background. **d**, Schematic depiction of DEPAF architecture. It uses a two-branch pipeline to process an original image, POI examples, and a detection sensitivity hyperparameter to output POI positions and a synthesized clean image with only the signals. FM, fitting map.

Here, we present DEep PAttern Fitting (DEPAF), a self-contrastive learning-based [34, 35] fluorescence microscopy signal detection framework featuring interference robustness *without* interference modeling, high generalizability, and unsupervised training within a compact, user-friendly design (Fig. 1b). Leveraging the inherent structural regularity of fluorescence microscopy signals, such as point spread functions (PSFs) [1, 36], periodic fluctuations [22, 37], and specific frequency distributions [8], DEPAF utilizes repetitive combinations of small data segments containing “patterns of interest” (POIs) to universally represent these diverse signals. By contrasting the consistency between different augmented views of the input data generated asynchronously via asymmetric model paths, DEPAF learns discriminative features determined solely by interference level rather than specific interference models. These features are embedded into a newly introduced subpixel-level dense representation scheme, ensuring strong discriminability between overlapping signals and thereby substantially improving the detectable signal density, even under conditions of strong noise and severe background interference with unknown types and parameters.

We demonstrated the generalizability and signal detection performance of DEPAF across a range of representative tasks involving diverse imaging modalities (Fig. 1c), requiring only the replacement of POI examples and the adjustment of a single detection sensitivity hyperparameter. In localization-based super-resolution microscopy, DEPAF enabled an order-of-magnitude improvement in 2D PSF detection rate under low-SNR conditions, even at a signal density of ∼30 PSFs/*µ*m^2^. This enhancement improved the temporal resolution of 2D single-molecule localization microscopy (SMLM) from several seconds to 76.8 ms without sacrificing spatial resolution, thereby enabling super-resolution imaging of dynamic minute changes in live cells at the cytoskeletal level with a spatial resolution of ∼30 nm. DEPAF also achieved high-robust-accuracy 3D SMLM reconstruction, supporting molecular blinking densities 10 times higher than conventional methods while requiring only 10.8% of the average photon count compared to typical STORM data labeled with Alexa647 [36]. In spatial transcriptomics imaging, DEPAF enables high-sensitivity detection of multiplexed error-robust fluorescence in situ hybridization (MERFISH) spot signals in strong background fluorescence. This results in a 97% increase in detection rate and an improvement of 0.12 in the correlation between detected RNA copy numbers and fragments per kilobase per million reads (FPKM) in a 3D MERFISH sample with strong background fluorescence. In low-SNR two-photon calcium imaging of neurons, DEPAF uses a single pipeline for denoising, background separation, weak spike inference, and active neuron segmentation. As an unsupervised source extraction-based segmentation method, DEPAF clearly outperforms other methods of the same type, increasing precision by 40% and recall by 43%, and is even competitive with top-performing supervised methods, while reducing the required training data and ground-truth labels by 2 and 1 orders of magnitude, respectively. DEPAF also allows for signalaware and accurate estimation of arbitrary inhomogeneous fluorescence backgrounds. Across simulated datasets with a wide range of background variation scales and SNR levels, the correlation between DEPAF-estimated and ground-truth backgrounds consistently exceeded 0.97 without introducing signal loss. These organic integrations demonstrate that DEPAF can serve as a practical framework, generally enhancing the signal detection performance under strong interference. With open-source code, flexible POI representation, and task-agnostic modeling, DEPAF is expected to bring broad benefits to related fields in the era of image-based multi-omics [38].

## Results

### DEPAF architecture

The architecture of DEPAF is illustrated in Fig. 1d. Our method takes an original image, POI examples, and a hyperparameter corresponding to detection sensitivity as inputs, and outputs the POI positions within the image along with a synthesized clean image that contains only the signals. The pipeline of our method comprises two branches. In the first branch, DEPAF treats each pixel as a fitting component and utilizes an image-to-image translation deep learning model with a U-net structure [39] to simultaneously (rather than sequentially) resolve all POIs in the original image while also denoising it. The result is represented as a 3D fitting map. The first and second dimensions of this map correspond to the likelihood of a POI’s presence at different pixels, while the third dimension corresponds to the likelihood of each POI’s existence at that pixel. The deep learning model in this branch is optimized by regularizing the fitting map to encourage sparsity and maximizing the similarity between the original image and the denoised image, where the denoised image is obtained by group-convolving the fitting map with the POI examples.

In the second branch, DEPAF uses an optimizable decoding module that leverages the principles of centroid-based discrete-to-continuous space transformation to decode the POI positions in continuous coordinate space from the fitting map. Then, the fitting map and the noise image, obtained by subtracting the denoised image from the original image, are combined to create a noisy fitting map, from which the amplitudes of the POIs are estimated. With the examples, positions and amplitudes of POIs, a synthesized image is generated. The decoding module is then optimized by maximizing the similarity between the synthesized and denoised image to remove noise-induced POI positions while retaining valid ones. After all the optimizations are complete, new image can be sequentially input into the fixed-parameter deep learning model and the decoding module for fast POI localization and image denoising.

### DEPAF benchmarking

We begin with a comprehensive benchmarking of DEPAF across four representative fluorescence microscopy signal processing tasks encompassing diverse data dimensions and modalities: 2D/3D SMLM, neuronal spike inference, and fluorescence background estimation (Fig. 2). The evaluation employed 43 simulated datasets, systematically covering a wide range of SNRs, signal densities, and background variation scales, enabling rigorous assessment of both accuracy and robustness. Compared to 17 leading methods in each domain, including supervised learning, simulation learning, and conventional statistical methods, DEPAF consistently matches or outperforms existing techniques across most settings. Notably, it remains stable under challenging conditions such as extremely low SNR or ultra-high signal density, where other methods often degrade. These results confirm DEPAF’s strong generalizability across tasks and resilience to interference. Specific details regarding each task are provided in the following sections.

**Fig. 2.**
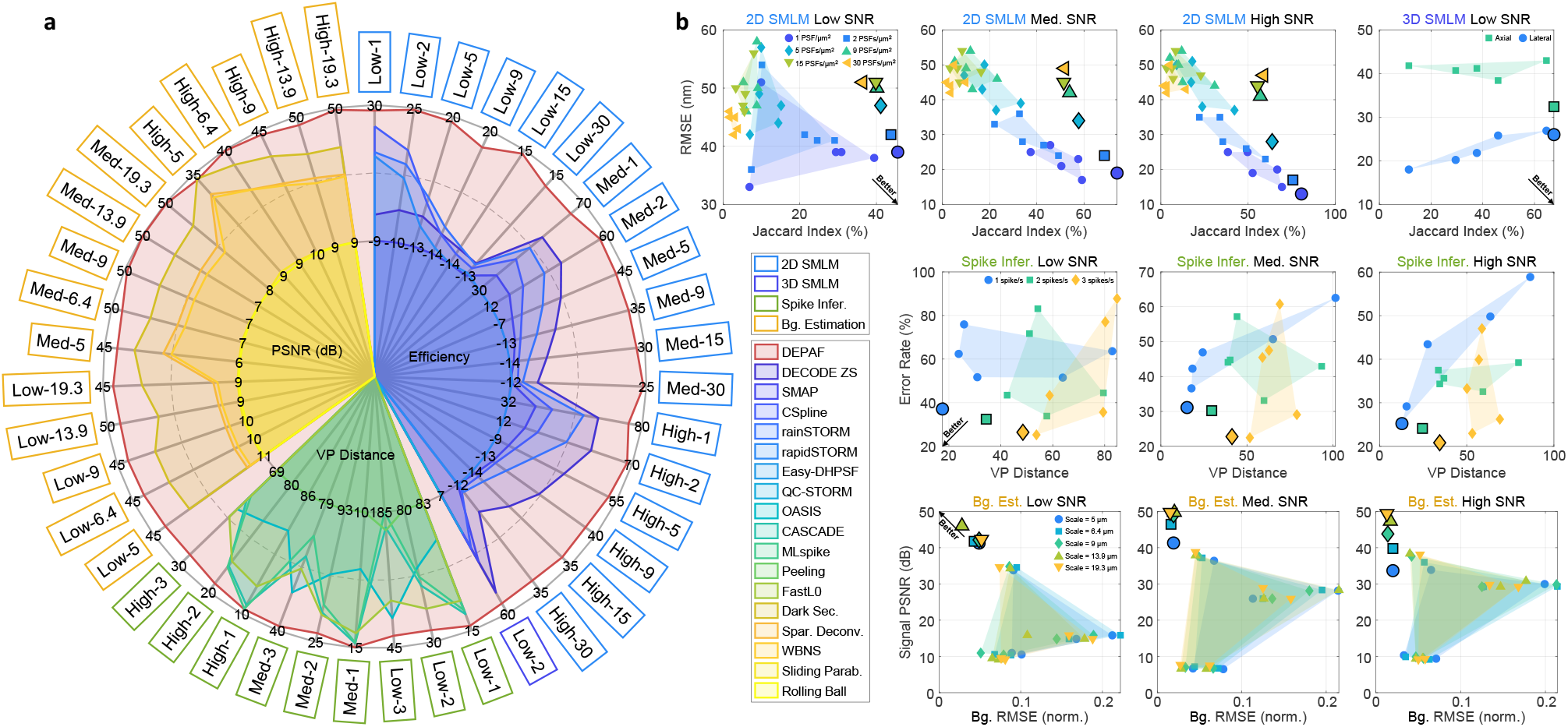
Benchmarking DEPAF on diverse and challenging simulated datasets. **a**, DEPAF outperforms a range of task-specific methods for 2D SMLM, 3D SMLM, spike inference, and background estimation across 43 simulated datasets. Performance was evaluated using summary metrics: Efficiency (higher is better) for SMLM, Victor–Purpura (VP) distance [40] (lower is better) for spike inference, and PSNR of background-removed signal images (higher is better) for background estimation. In dataset names, “Low”, “Med”(medium), and “High” indicate SNR levels; numbers denote signal density (e.g., PSF density or spike rate) or background variation scale. Infer., inference; Bg., background; ZS, zero-shot testing; Sec., sectioning; Spar., sparsity; Deconv., deconvolution; Parab., paraboloid. **b**, Paired trade-off metrics provide a detailed comparison, where small markers denote baseline methods and bold-edged large markers indicate DEPAF. DEPAF consistently lies on the Pareto front, highlighting its robust and generalizable performance. Med., medium; Infer., inference; Bg., background; Est., estimation; Norm., normalized.

### Millisecond-level dynamic 2D SMLM and high-robust-accuracy 3D SMLM via DEPAF

Diffraction causes a lens-based microscope to image a point light source as a PSF with a width of 200–300 nanometers [1], making structures smaller than this scale appear blurred and difficult to distinguish. SMLM techniques, such as STORM [41], dSTORM [42], PAINT [43], and PALM [44], are dedicated to addressing this issue by randomly activating fluorescent molecules within the samples, recording their “blinking” behavior in numerous camera frames, and then using localization algorithms to precisely localize the nanoscale position of each PSF for super-resolution imaging rendering. These techniques have transformed biological imaging, enabling nanoscale observation of subcellular structures.

Temporal resolution is inherently limited in SMLM by the sequential activation of fluorescent molecules and the requirement for capturing numerous frames [5]. Conventional SMLM typically achieve temporal resolution on the order of minutes, with some reports of tens of seconds [45, 46]. Solutions include: (1) allowing a higher density of PSFs per frame, (2) reducing the activation time of fluorescent molecules, and (3) increasing the imaging frame rate. This requires the localization algorithms of SMLM to accurately detect and localize highly overlapping PSFs in strong noise caused by fast frame rates.

To quantitatively test the performance of DEPAF in 2D SMLM, we set a calibrated PSF image as the POI example and evaluated it on 18 simulated 2D SMLM datasets, which spanned a wide range of SNR levels and high PSF densities. Compared to SMAP [15], the top-performing conventional statistical method in a public SMLM challenge [36], and DECODE [4], a simulation-based method that outperformed all existing methods across all 12 datasets in the same public challenge, DEPAF showed clear performance improvements in localizing ultra-highdensity PSFs and maintained exceptional robustness under low SNR imaging conditions (Fig. 3a and Extended Data Fig. 2). For instance, in the most challenging dataset with a low SNR and an ultra-high density of 30 PSFs/*µ*m^2^, DEPAF increases the Jaccard index, which measures the detection rate, from 3.87% with DECODE and 1.64% with SMAP to 36.41% (Extended Data Fig. 2a), achieving an order-of-magnitude improvement. This demonstrates DEPAF’s strong ability to address the temporal resolution limitations in SMLM.

**Fig. 3.**
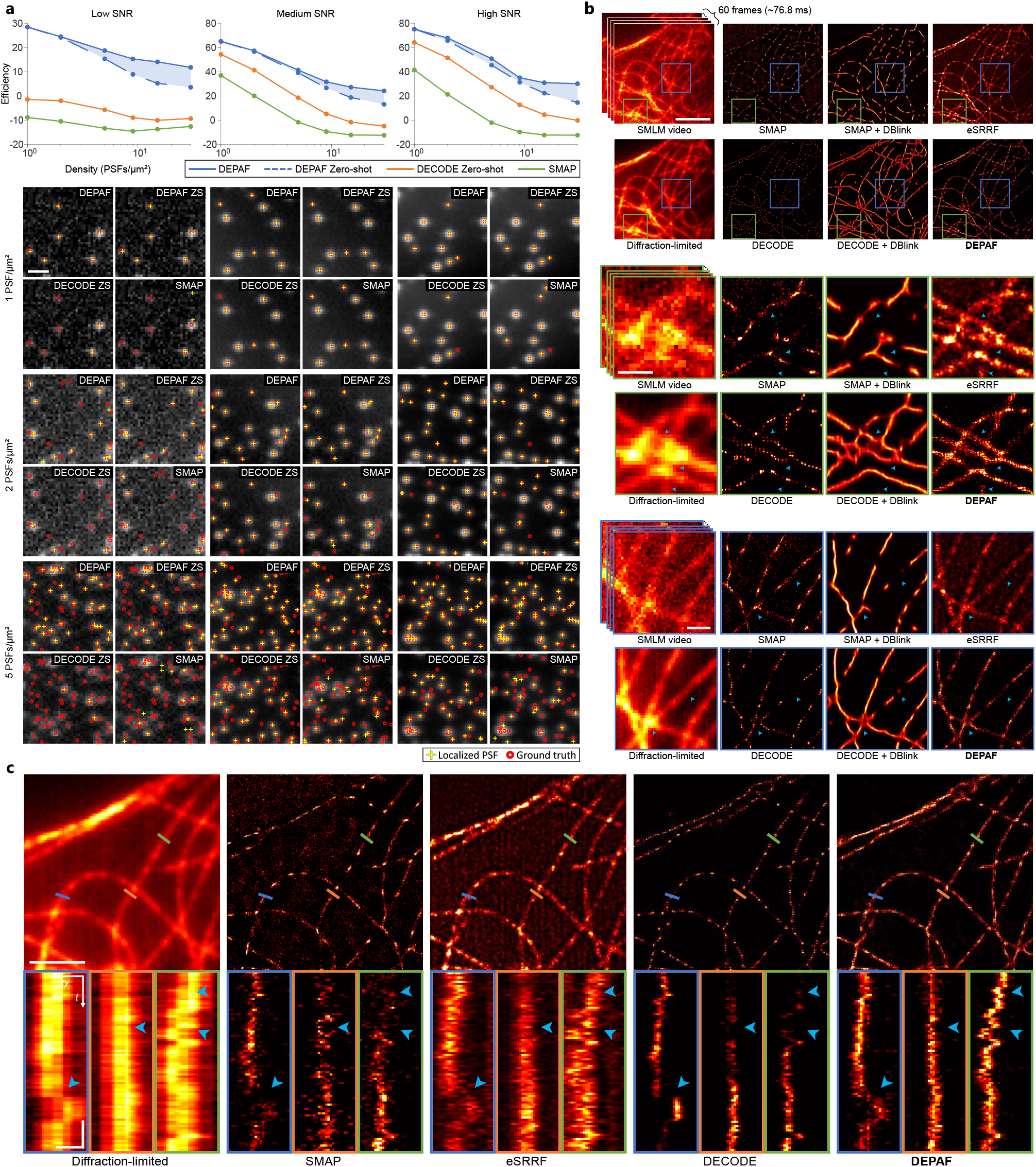
DEPAF outperforms other methods in all 18 simulated SMLM datasets with a wide range of SNR levels and high PSF densities and achieves millisecond-level dynamic 2D SMLM. **a**, The line charts compare the *efficiency* metrics of different methods across different SNR levels and PSF densities (log scale) in 18 simulated datasets (see Extended Data Fig. 2 for further details). Each column of subplots below visualizes the localization results at PSF densities of 1, 2, and 5 PSFs/*µ*m^2^ for the corresponding low, medium, and high SNR levels. Scale bar, 1 *µ*m. ZS, zero-shot testing. **b**, Top: super-resolution reconstruction results of different methods based on SMLM videos captured with a high-speed camera at a frame rate of 1.28 ms/frame and the dSTORM technique with high blinking density. Scale bar, 5 *µ*m. Middle and bottom: magnified details of the reconstruction results. Blue arrows indicate specific regions to highlight differences in resolution and structural details among the methods. Scale bar, 1 *µ*m. **c**, The subplots below display the time-lapse images along the line segment in the top images within the reconstructed super-resolution video. Differences in resolution and structural details between the methods are indicated by blue arrows in specific areas. Vertical scale bar, 1 s. Horizontal scale bar, 300 nm.

Using the dSTORM technique [42], we performed SMLM imaging on a live-cell sample to capture cytoskeletal fluctuations with high blinking density at a frame rate of ∼781 frames per second (FPS) (Fig. 3b) (Supplementary Note 3). Subsequently, using a calibrated 2D PSF as the POI example, DEPAF achieves the rendering of super-resolution videos with a temporal resolution of ∼76.8 ms ( ∼13 FPS) and a spatial resolution of ∼30 nm (Fig. 3b,c and Supplementary Video 1). The spatial resolution is quantified using decorrelation analysis [47] (Supplementary Fig. 1). This enables the observation of rapid, minute changes in subcellular structures at the cytoskeletal level. In comparison, methods such as SMAP, which relies on sparse signal conditions, DECODE, which have been reported to achieve a temporal resolution of a few seconds [4], and eSRRF, which can generate one super-resolution volume per second but compromises spatial resolution to ∼70 nm [37], as well as DBlink, which achieves millisecond-level resolution but is limited by interpolation-induced artifacts [5], are all unable to achieve high-quality super-resolution reconstructions under such challenging imaging conditions (Supplementary Video 1).

PSF engineering can employ special optical components or modulation methods to modify the PSF shape produced by emitters at different *z*-axis positions [48]. This allows for the estimation of the *z*-axis positions of fluorescent molecules based on their PSF shapes in 2D SMLM frames, enabling 3D SMLM [36, 49]. These engineered PSFs often show non-orthogonality and greater overlap in the frames due to their larger area, which raises the performance demands on localization algorithms.

Using 151 calibrated double helix PSFs [50] as the POI examples, we evaluated DEPAF on a simulated 3D SMLM benchmarking dataset, where the average PSF density per frame was 10 times higher than normal, and the average photon count was only 10.8% of the typical data acquired from Alexa647-labeled STORM samples [36] (Extended Data Fig. 3a–c). The results show that DEPAF can effectively handle these challenging conditions and achieve high-quality reconstruction, outperforming leading statistical and simulation-based methods tailored for this task (Extended Data Fig. 3d,e).

### High-sensitivity 3D MERFISH spot signal detection using DEPAF

Single-molecule fluorescence in situ hybridization (smFISH) utilizes fluorescence labeling, imaging, and algorithm-based spot signal detection to precisely localize specific RNAs with high resolution [51, 52]. This makes smFISH a powerful tool for studying RNA copy number and spatial organization. With technological advancements, more sophisticated techniques, such as MERFISH, have further expanded the detection capabilities, allowing for the simultaneous analysis of hundreds to thousands of RNA species [2]. However, these advancements also introduce challenges, such as the need for additional rounds of hybridization, increased non-specific hybridization, and higher labeling densities [53]. These factors can result in stronger fluorescence backgrounds [7] and greater signal overlap [54] in the images. Current spot signal localization algorithms still struggle to effectively tackle such complex data.

To demonstrate DEPAF’s ability to overcome these challenges, using a fitted Gaussian PSF image as the POI example, we applied it to a 3D *z*-stack MERFISH data obtained from a fetal liver tissue sample. This data is characterized by strong fluorescence backgrounds, high-density labeling, and a large FOV of 165.7 *×* 165.7 *×* 9 *µ*m^3^ [54]. In addition to the typical challenges of conventional 2D MERFISH data, *z*-stack imaging introduces further complexities, such as signal crosstalk between *z*-layers, PSF defocusing, *z*-axis drift, and more pronounced fluorescence intensity fluctuations, which pose additional difficulties to the spot signal detection algorithms.

Our results show that, compared to the 68,294 RNA molecules detected by the official tool [54] based on conventional statistical method, DEPAF substantially increases the detection number to 134,560, resulting in a 97% improvement in detection rate (Fig. 4a,b). To verify the reliability of the detection results, we employed a standard evaluation method by comparing the detected RNA copy numbers with the abundance determined by bulk sequencing in FPKM on the same cell line [2]. The results demonstrate that DEPAF further improves the correlation between RNA copy numbers and FPKM by 0.12 compared to the official tool (Fig. 4c), reaching a usable threshold of 0.7 [54]. Additionally, DEPAF reduces the average miss rate of bits from 23.1% to 14.3% while slightly increasing the average false positive rate from 0.8% to 1.7% (Fig. 4d).

**Fig. 4.**
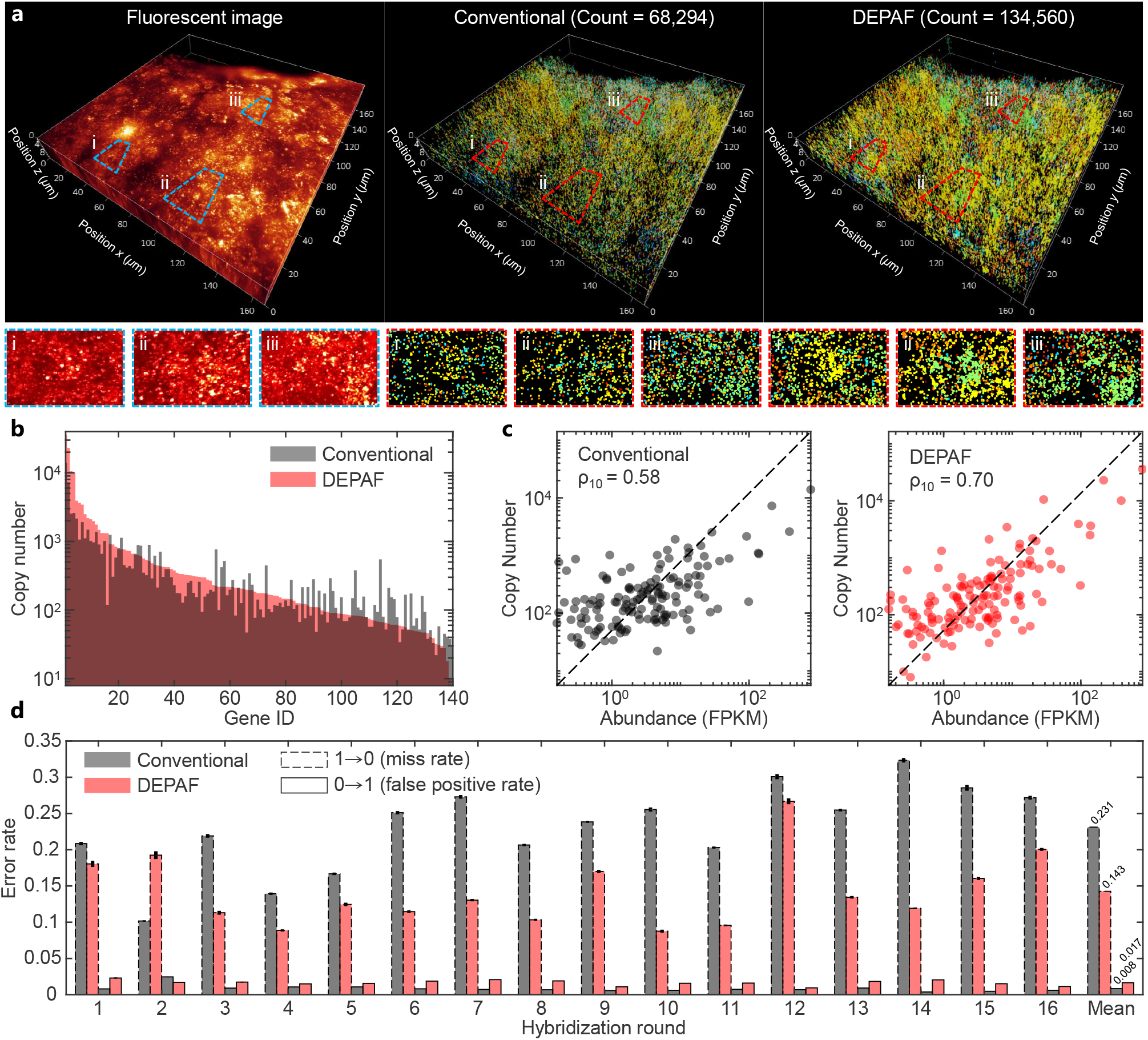
DEPAF doubles the detection rate of MERFISH spot signals even under complex imaging conditions with strong backgrounds, high labeling density, and a large FOV. **a**, In a 3D MERFISH data labeling 140 RNA species acquired in challenging conditions (left), the number of RNA molecules detected by DEPAF (right, 134,560 counts) is considerably higher than that detected by the conventional method (middle, 68,294 counts). The zoomed-in view at the bottom provides detailed visualization of the image and detection results. The 3D fluorescence image is shown as a maximum intensity projection created by combining images from different hybridization rounds after alignment and bleaching correction. The subplots below display maximum intensity projections of different slices derived from this image. Different colors represent different RNA species. **b**, Scatter plots compare the copy numbers detected by the conventional method (left, black) and DEPAF (right, red) with the abundance measured by bulk sequencing. The Pearson correlation coefficients between the log_10_-transformed copy numbers and abundances (*ρ*_10_) are 0.70 and 0.58, with a *P* value of 5 *×* 10^−22^ and 5 *×* 10^−14^, respectively. **c**, The bar plot shows the RNA copy numbers detected by DEPAF (red) and the conventional method (black) across different RNA species (log_10_ scale). Gene IDs are sorted in descending order based on the RNA copy numbers detected by DEPAF. **d**, The bar plot displays the miss rates (1 *→* 0) and false positive rates (0 *→* 1) calculated based on error correction with MHD4 coding [2] across multiple hybridization rounds for DEPAF (red) and the conventional method (black). Error bars represent standard error of the mean (SEM, *n* = 16 *z*-layers).

### Low-SNR two-photon calcium imaging data processing with DEPAF

By using genetically encoded calcium indicators and two-photon microscopy, we can transform neuronal action potentials (spikes) into visible fluorescent fluctuations [3]. This allows us to efficiently conduct large-scale optical recordings of neuronal activity in various animal models and associate these activities with neural functions, thereby substantially enhancing our understanding of how local neural networks process information.

Extracting neural activity from two-photon calcium imaging videos involves two key tasks: (1) reconstructing spikes and estimating their positions in the temporal domain [21–24, 29], and (2) extracting and segmenting active neurons in the spatial domain [25–28]. However, the imaging data often suffers from significant noise interference, particularly in high temporal resolution analyses [6], and exhibits spatiotemporally inhomogeneous backgrounds [22]. Moreover, a single spike manifests as rapid rises followed by slow decays, which may lead to spike overlap [55]. These factors pose challenges to spike inference and active neuron segmentation.

Leveraging DEPAF’s strong performance under low-SNR and high-density spike inference benchmarks (Extended Data Fig. 4), we applied it to video J123 from the CaImAn dataset [25], a representative low-SNR two-photon calcium imaging data of hippocampal neurons (Fig. 5a). The results show that, considering the data along the time axis for each pixel in the two-photon calcium imaging videos as a one-dimensional (1D) image and using a fitted spike signal as the POI example, DEPAF can localize overlapping and weak spikes in significant noise (Fig. 5b,(i)–(iv)). Meanwhile, benefiting from DEPAF’s accurate estimation of POI positions and amplitudes (Extended Data Fig. 5) and through its image synthesis function (Fig. 1d), noise-free signals separated from the background can be generated (Fig. 5b,c and Extended Data Fig. 6a,b). After reshaping these noise-free signals into 3D videos, denoised two-photon calcium imaging videos that remove the background and retain only active neuron signals can be obtained (Fig. 5b and Extended Data Fig. 6c).

**Fig. 5.**
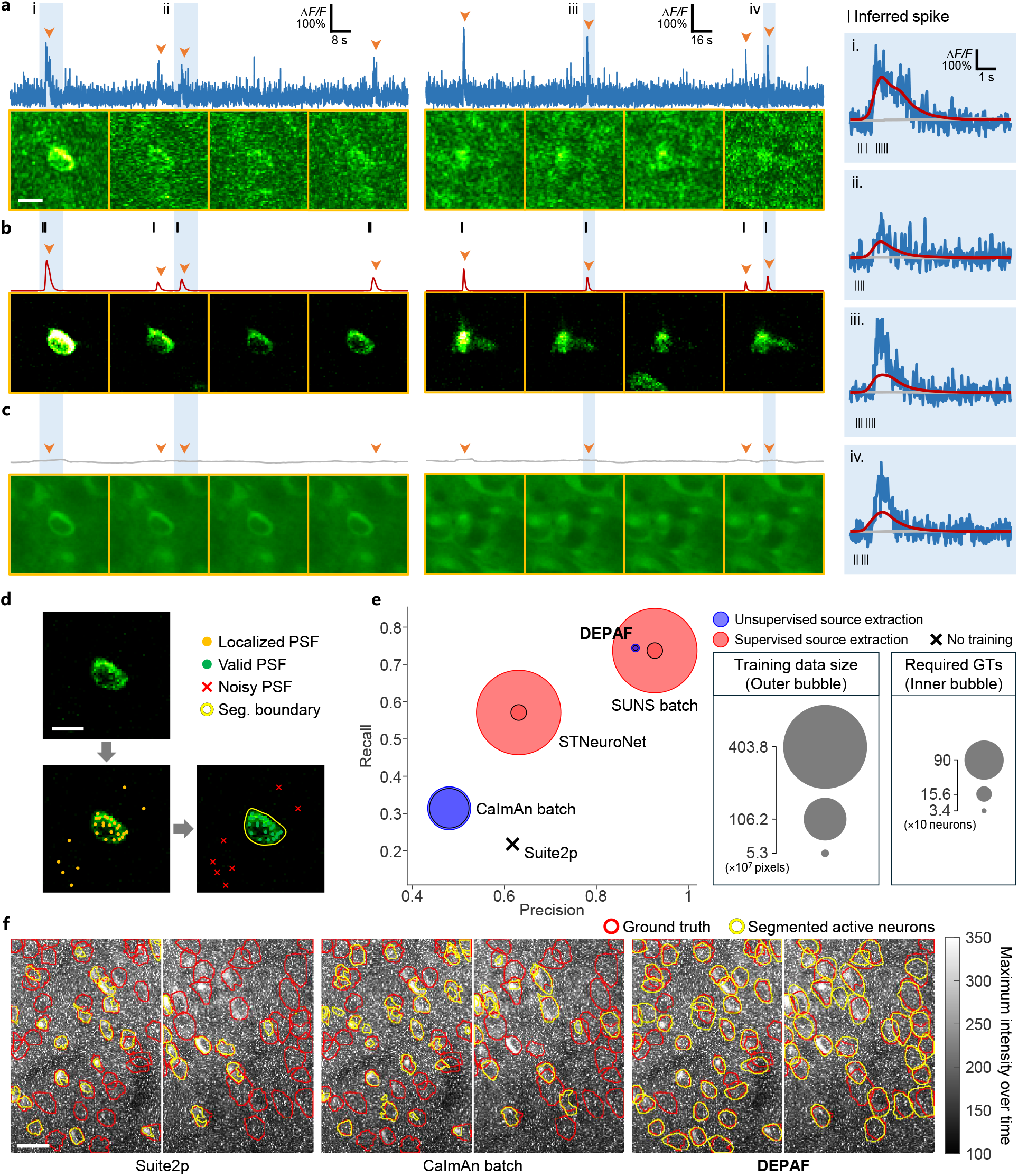
DEPAF enables one-pipeline processing for denoising, background separation, weak spike inference, and data-efficient active neuron segmentation in low-SNR two-photon calcium imaging data. **a**, Fluorescence traces of two neurons from the low-SNR two-photon calcium imaging data. Frames marked with orange arrows are shown below the traces. Δ*F* represents the change in fluorescence relative to the baseline fluorescence *F*_0_, which is approximated by the average of the entire trace. Scale bar, 9 *µ*m. **b**, Fluorescence traces and frames after DEPAF denoising, retaining only signals from active neurons. Spike positions are simultaneously estimated with DEPAF and indicated by black vertical lines (variations in thickness are due to overlaps). **c**, Fluorescence traces and frames of the spatiotemporally inhomogeneous background estimated by DEPAF. **i**–**iv**, Zoomed-in examples of calcium transients in raw data (blue), denoised data (red), estimated background (gray), and estimated spike positions (black vertical lines). **d**, The process of active neuron segmentation using the denoised data. DEPAF detects potential signals using a standard Gaussian PSF as the POI example, filters valid PSFs via DBSCAN clustering, and generates segmentation boundaries using cubic spline curves based on the valid PSFs. Scale bar, 9 *µ*m. Seg., segmentation. **e**, Segmentation performance of different methods, along with their training data size and ground-truth labels used. GTs, ground-truth labels. **f**, Example segmentation results of all *unsupervised* source extraction-based methods from the second and fourth quadrants of video J123, overlaid on the maximum intensity projection across the time dimension. Scale bar, 20 *µ*m.

Based on the denoised videos (Supplementary Video 2), DEPAF can further detect the ultra-high-density PSFs that constitute the active neuron regions in the spatial domain using a standard Gaussian PSF image as the POI example. After localization, active neurons are extracted and segmented by performing DBSCAN clustering [56] and boundary generation on the localized PSFs (Fig. 5d). Comparative experiments on active neuron segmentation demonstrate that DEPAF, a method based on unsupervised source extraction for identifying the spatial footprints of active neurons, substantially outperforms other methods that likewise do not require groundtruth supervision for source extraction, such as Suite2p [26] and CaImAn batch [25]. In particular, DEPAF achieves a 40% increase in precision and a 43% increase in recall compared to CaImAn batch (Fig. 5e,f). To further benchmark the performance of DEPAF, we also compared it against supervised methods (Supplementary Fig. 2). The results show that DEPAF still outperforms STNeuroNet [28] and even performs competitively with the top-performing supervised method SUNS batch [27], despite requiring two orders of magnitude less training data and one order of magnitude fewer ground-truth labels (Fig. 5e).

### Signal-aware estimation of arbitrary inhomogeneous fluorescence background enabled by DEPAF

In fluorescence microscopy images, “background” refers to components other than the signals and noise. These components reduce image quality and affect the sensitivity and accuracy of signal detection, necessitating their removal. Assuming the background is homogeneous [15, 16] oversimplifies real imaging environments. Typical specimens, such as cells or tissue slices, contain various components at different spatial scales that may fluoresce spontaneously. Additionally, fluorescent markers might bind non-specifically to other components. Thus, the actual background is an inhomogeneous mixture of various spatial frequencies [4, 8].

For DEPAF, the background can likewise be treated as a special type of “signal” to be detected. Thus, accurately estimating arbitrary inhomogeneous backgrounds hinges on how well the background POI example can broadly represent real background structures. Previous studies have shown that background distributions are concentrated in low frequencies [8], meaning the background POI example should capture low-frequency information. Additionally, since the background is distinct from both signals and noise, the background POI example must be uncorrelated with them to prevent DEPAF from mistakenly fitting it to either. Therefore, the background POI example should be designed to be orthogonal to both. To satisfy these requirements, we designed the background POI example as a homogeneous patch of a certain size, essentially a small image of constant value. This offers the following benefits: (1) Its size limits its ability to fit high-frequency information, making it suitable for low-frequency representation. (2) Due to the periodic nature of signals and the randomness of noise, this homogeneous patch maintains broad orthogonality to both. These features align well with the requirements. Thus, benefiting from DEPAF’s capability to fit multiple POI examples simultaneously, by using this background POI example as an additional one alongside signal POI examples, DEPAF can optimize background estimation and signal detection simultaneously, thereby achieving signal-aware estimation of fluorescence backgrounds.

Experimental results on 15 simulated datasets demonstrate that DEPAF estimates arbitrary inhomogeneous backgrounds with correlation to the ground truth consistently above 0.97 across a wide range of background variation scales and SNR levels. Compared to Dark Sectioning and WBNS, DEPAF yields background estimates better separated from the signal, thereby avoiding signal loss in the background-removed images (Fig. 6a,b and Extended Data Fig. 7). It also exhibits excellent performance on experimental data (Fig. 5c, Fig. 6c,d, and Extended Data Fig. 6c).

**Fig. 6.**
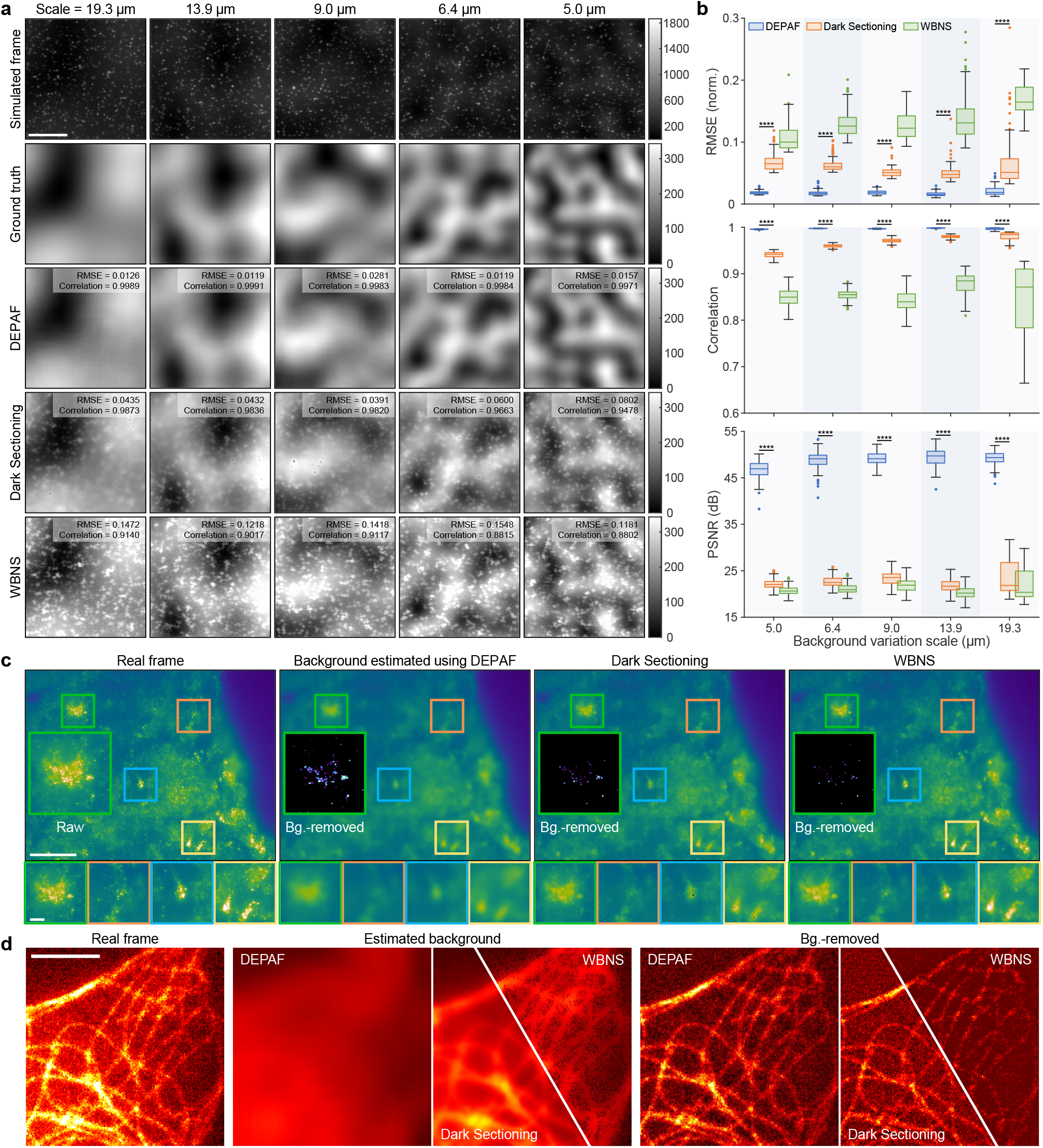
DEPAF accurately estimates inhomogeneous backgrounds without compromising signals. **a**, Performance comparison of DEPAF, Dark Sectioning, and WBNS, under medium SNR level and different background variation scales (see Extended Data Fig. 7 for low- and high-SNR results). First row: simulated fluorescence frames. Second row: ground-truth back-grounds. Rows three to five: backgrounds estimated by DEPAF, Dark Sectioning, and WBNS, respectively. RMSE and Pearson correlation coefficients with ground truth are shown in the top right of each image. Scale bar, 10 *µ*m. **b**, Quantitative comparison of RMSE, Pearson correlation coefficients, and PSNR of background-removed images across background variation scales. \*\*\*\**P <* 0.0001 for all comparisons. *P* values are calculated with the two-sided *t*-test. Norm., normalized. **c**, Background estimation on a real MERFISH frame. Left to right: raw image and backgrounds estimated by DEPAF, Dark Sectioning, and WBNS. Scale bar, 30 *µ*m. Insets show background-removed results, highlighting DEPAF’s ability to suppress background without distorting signals. Bottom panels show zoomed-in views of estimated backgrounds. Scale bar, 5 *µ*m. Bg., background. **d**, Background estimation on a SMLM frame of cellular microtubules. Shown are the raw image, estimated backgrounds, and background-removed results from DEPAF, Dark Sectioning, and WBNS. DEPAF preserves fine filamentous structures while suppressing uneven background more effectively than other methods. Scale bar, 5*µ*m. Bg., background.

## Discussion

In this work, we present DEPAF, an interference-resilient framework for fluorescence microscopy signal detection that eliminates the need for interference modeling. This feature further imparts generalizability to the method, which is tested across multiple tasks that essentially encompass different representative signal types: To demonstrate its effectiveness in detecting single-shape signals, we applied it to 2D SMLM, achieving millisecond-level dynamic super-resolution imaging; To showcase its performance in detecting multi-shape signals, we employed it in 3D SMLM, achieving high-robust-accuracy 3D super-resolution imaging; To test its ability to handle signals without precise shape calibration, we used it in MERFISH analysis, supporting high-sensitivity 3D MERFISH spot signal detection in strong background fluorescence; To further demonstrate its capability in processing cross-dimensional data and various potential applications, such as 1D signal detection, blind denoising, and geometric segmentation through point localization, we applied it to low-SNR two-photon calcium imaging video processing, achieving denoising, background separation, weak spike inference, and data-efficient active neuron segmentation in a single pipeline; Lastly, to demonstrate its ability to detect signals under broader priors unrelated to signal shape, we applied it to fluorescence background estimation, enabling accurate estimation of arbitrary inhomogeneous backgrounds. These cases suggest that DEPAF holds considerable potential for further development.

DEPAF is user-friendly, as the adaptation for different tasks can be achieved by simply replacing the POI examples and adjusting a hyperparameter related to detection sensitivity. This is possible due to the structural and regular features of fluorescence microscopy signals can be broadly and uniformly represented through POI, and the model architecture used strikes a good balance between specificity and generality. For instance, it focuses only on interference level rather than the exact type and parameters of interference; It utilizes general priors aligned with physical intuition to decode fitting results from discrete pixel space to continuous space; It utilizes contrast relationships for data processing without introducing additional manually set parameters or external ground truth. These make DEPAF lower the technical barriers to accessing high-performance fluorescence microscopy signal detection. By simplifying operations, it requires only minimal and intuitive input, eliminating the need for users to collect labor-intensive labeled datasets or parameterize physics-based imaging models that rely on specialized expertise.

The framework of DEPAF is scalable. As a model-agnostic solution, the U-net used in DEPAF serves as a simple end-to-end baseline, which can be easily extended to more advanced network architectures in the future. DEPAF also introduces an important perspective: various fluorescence microscopy signals can be represented through different POI examples. This means DEPAF can expand into a wider range of applications by incorporating more well-designed POI examples. Building on this concept, as demonstrated in tasks such as 3D SMLM and fluorescence background estimation, DEPAF supports co-optimization using multiple POI examples to simultaneously detect multiple types of signals. Therefore, its scalable and integrable architecture positions it as a potential unified model to address the lack of foundational frameworks in the field.

In conclusion, DEPAF has transformed the performance benchmarks of current fluorescence microscopy signal detection methods while maintaining its generalizability and scalability. We envision it as a standardized, high-performance signal detection framework applicable across cellular, subcellular, and molecular scales, facilitating the advancement of emerging techniques such as image-based multi-omics [38], localization-based live-cell super-resolution imaging, and their integration.

## Methods

### DEPAF pipeline

Our pipeline consists of two branches. In the first branch (Extended Data Fig. 1a), a deep learning model ℳ_net_ is initialized and used to transform the original image *I*_ori_ into a fitting map *I*_fit_, which encodes the POI positions and amplitudes. This map is further group-convolved with the POI examples to produce a denoised image *I*_dns_. The ℳ_net_ is optimized by maximizing the *L*_2_ similarity between *I*_ori_ and *I*_dns_ with the *L*_1_ regularization of *I*_fit_. Once ℳ_net_ is optimized, in the second branch (Extended Data Fig. 1b), an optimizable decoding module ℳ_decode_ is initialized and utilized to decode the POI positions from *I*_fit_. By subtracting *I*_dns_ from *I*_ori_, a noise image *I*_ns_ is obtained. This noise image is then merged with *Î*_fit_, the sum of *I*_fit_ in the channel dimension, to produce a noisy fitting map *Ĩ*_fit_. The POI amplitudes are estimated based on *Ĩ*_fit_. Subsequently, the estimated POI amplitudes are combined with the known POI examples and the obtained POI positions to generate a synthesized image *I*_syn_. The ℳ_decode_ is optimized by maximizing the *L*_1_ similarity between *I*_syn_ and *I*_dns_. After the optimizations are complete, the parameter-fixed ℳ_net_ and ℳ_decode_ can be utilized for POI detection on new images. The detailed process is presented below.

#### POI example construction

DEPAF applies min-max normalization to each of the *C* pre-acquired POI images 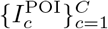, scaling their pixel intensities to [0, 1]. The normalized images are then stacked along the channel dimension to form the POI examples *P* ∈ ℝ^*W ×H×C*^, where 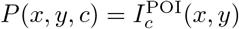.

#### First branch

In this branch, the ℳ_net_, a full convolutional U-net neural network [39] (Extended Data Fig. 1c), is used to simultaneously fit all the POIs in *I*_ori_ and denoise the *I*_ori_. The loss function for optimizing ℳ_net_ is:

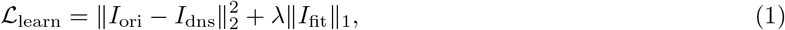

where *I*_ori_ is the original image input to the model, *I*_fit_ is the fitting map output from the model, *λ* is the *L*_1_ regularization strength, and *I*_dns_ is the denoised image calculated by:

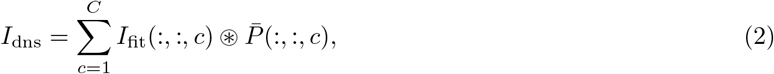

where “⊛” denotes a convolution operation, “:” denotes all indices along a dimension, 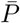 is obtained by dividing the POI examples by the mean of the total pixel sums across all channels. The loss function ℒ_learn_ comprises of an *L*_2_ similarity term and an *L*_1_ regularization term. The *L*_2_ similarity term enforces similarity between *I*_dns_ and *I*_ori_ for reconstruction, while the *L*_1_ regularization term promotes the sparsity of *I*_fit_ for denoising. The regularization strength *λ*, as a adjustable hyperparameter, can therefore control the denoising strength and sensitivity of POI detection.

#### Second branch

In this branch, the ℳ_decode_, an optimizable decoding module, is used to decode the POI positions from *I*_fit_. Since POI examples also share similarities with the noise in *I*_ori_ (not just with the POIs), *I*_fit_ may also contain fitting values produced by noise rather than by POIs. These noise fitting values typically do not exceed the noise level. Therefore, we can use a threshold *θ* to filter them out. However, since the noise level is difficult to estimate [57], an iterative optimization method is employed here to robustly address this issue. Specifically, We first sum *I*_fit_ along the channel dimension to obtain *Î*_fit_. Then we initialize a *θ* to be optimized within the pixel value range of *Î*_fit_. The fitting map after filtering out the noise fitting values is obtained by:

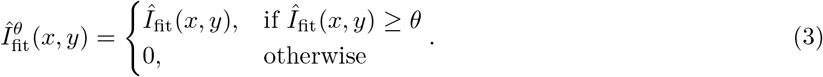

To further partition the regions of interest (ROIs) containing an individual POI in 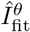, the following two priors (Extended Data Fig. 1d) are considered.

- **Shape prior:** Due to the discrete nature and inherent size of pixels, a point in continuous coordinate space is represented as 1 to 4 pixels within a 2 *×* 2 pixel block in discrete pixel space under a sparsity constraint, which corresponds to the shape of each ROI.
- **Priority prior:** For ROIs containing 1 to 4 non-zero pixels, the range of points in continuous coordinate space they can represent increases sequentially. Following classical probability theory, the likelihood of their occurrence also increases accordingly. Their priority for partitioning is thus in ascending order.

Therefore, the pseudo-code of an algorithm (Extended Data Fig. 1e) for using the above priors to partition ROIs is as Algorithm 1.

##### Algorithm 1

ROI Partitioning

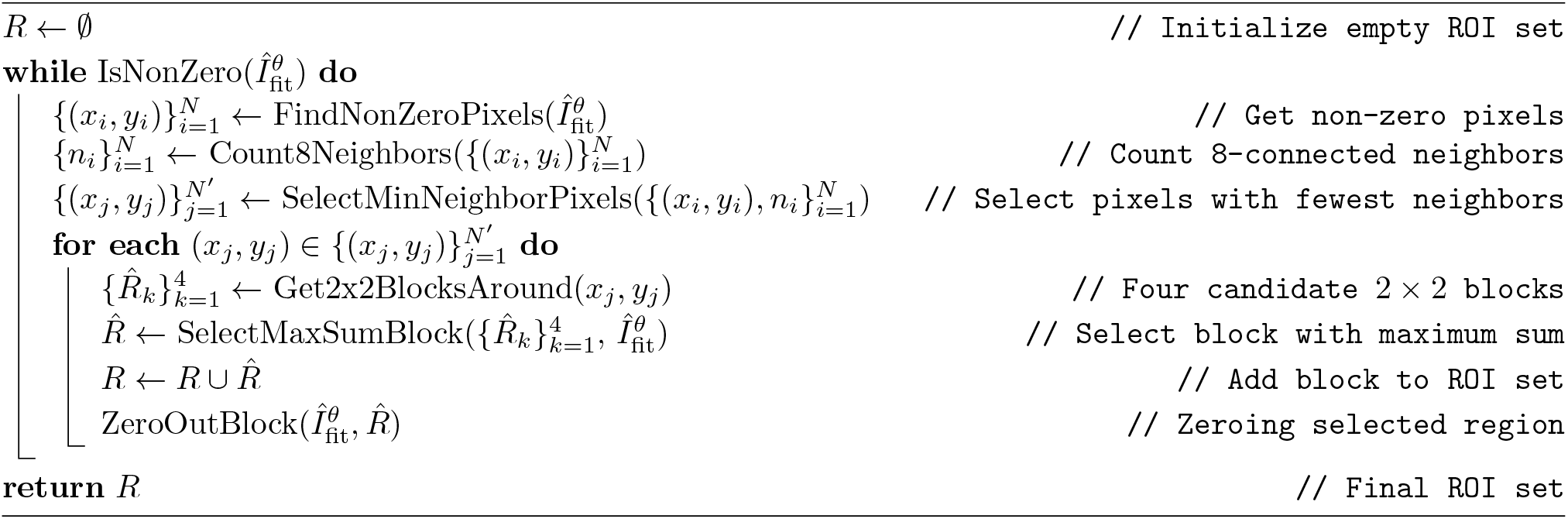

After ROI partitioning, the POI position 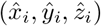 of *i*-th ROI 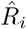 is estimated by calculating the center of mass:

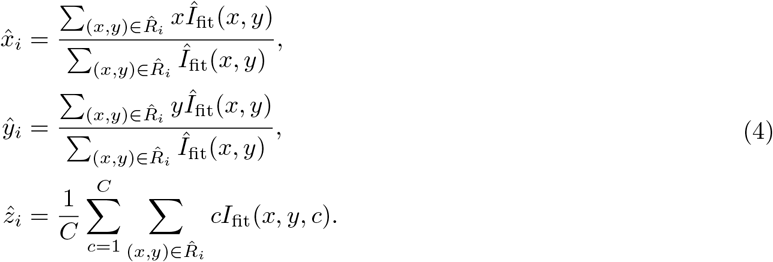

To further obtain the POI amplitudes, we first derive the noise image *I*_ns_ by subtracting the denoised image *I*_dns_ from the original image *I*_ori_. We then normalize *I*_ns_ using the maximum value of 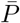 to ensure it is on the same scale as *Î*_fit_. These are then synthesized into a noisy fitting map by:

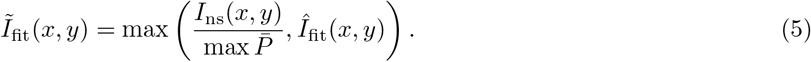

The purpose of this step is to use noise to contaminate the fitting values below the noise level while preserving those above the noise level. Afterward, the POI amplitude *A*_*i*_ of *i*-th ROI 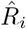 is estimated by calculating the maximum value after convolving each ROI with the POI example corresponding to its *z*-axis position:

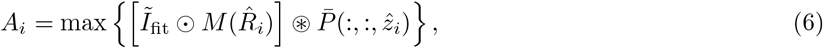

where “*⊙*” represents element-wise multiplication, *M* (·) is a function that outputs a mask where the pixels within the input set are 1 and the rest are 0. The sub-pixel values of the POI example at 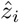 are obtained using interpolation. Based on the known POI examples and the obtained POI positions and amplitudes, a synthesized image *I*_syn_ can be generated by:

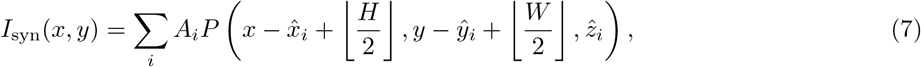

where *H* and *W* are the height and width of the POI examples, respectively. The sub-pixel values of the POI example at 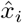 and *ŷ*_*i*_ are also obtained through interpolation.

Building upon the above construction, a critical trade-off emerges: choosing a too small *θ* that is below the noise level will result in the noise fitting values in *Î*_fit_ being retained. This leads to the generation of multiple POIs in *I*_syn_ with amplitudes derived from the noise in *Ĩ*_fit_. Due to the randomness and instability of noise, these noise-induced POIs cannot match the illusionary POIs in *I*_dns_, which are obtained by convolving the noise fitting values with the POI examples. Therefore, *I*_syn_ will deviate from *I*_dns_. Conversely, choosing a too large *θ* that is above the noise level will lead to the erroneous filtering of non-noise fitting values in *Î*_fit_. This results in fewer non-noise-induced POIs being generated in *I*_syn_ compared to those in *I*_dns_, similarly causing *I*_syn_ to deviate from *I*_dns_. Therefore, the *θ* can be optimized by maximizing the similarity between *I*_dns_ and *I*_syn_, i.e., minimizing the loss function:

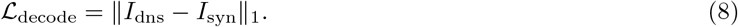

Due to the non-differentiability of ℒ_decode_ with respect to *θ*, a gradient-free optimization method is used here. Specifically, we start with Bayesian optimization [58, 59] to quickly estimate the shape of the ℒ_decode_-*θ* curve and find a suboptimal *θ*. After that, we perform a grid search method [60] centered around this suboptimal *θ* to find the optimal *θ*.

### Enhancement techniques

Three enhancement techniques are employed in DEPAF: (1) center-aligned upsampling to improve POI localization accuracy (Supplementary Fig. 3), (2) inhomogeneous background learning to enhance interference resistance (Supplementary Fig. 4), and (3) POI diffusion to promote model convergence in multi-POI detection scenarios (Supplementary Fig. 5).

#### Center-aligned upsampling

The inherent pixel size limits DEPAF’s ability to distinguish overlapping POIs and accurately estimate their positions using pixel-weighted centroids. To address this limitation, we reduce the pixel size by applying the same upsampling factor to both the POI examples and the original images at the start of our pipeline. To prevent localization bias, this process must ensure that the center position of the POI example remains consistent before and after upsampling. A detailed derivation and additional explanations of this method are presented in Supplementary Note 1.

#### Inhomogeneous background learning

We add a homogeneous patch as the background POI example *P*_bg_ and an output channel of ℳ_net_ to generate a background fitting map *I*_fitBG_. This method enables the model to learn arbitrary inhomogeneous backgrounds while detecting signals. When this technique is applied, the loss function to optimize ℳ_net_ becomes:

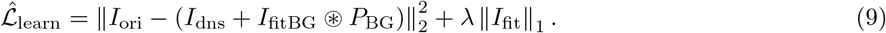

Note that since the background does not exhibit natural sparsity, *I*_fitBG_ is not constrained by *L*_1_ regularization.

#### POI diffusion

When our method is applied to multi-POI detection tasks, the number of POI examples determines the channel number of the output *I*_fit_ in ℳ_net_ (Equation 2). Therefore, as the number of POI examples increases, the search space of *I*_fit_ also expands, potentially affecting the convergence stability of ℳ_net_. To address this, we introduce the POI diffusion technique. This technique first employs bisection to divide all the POI examples into two groups. Within each group, mean pooling is used to merge the POIs into the central channel of that group. In the final convolutional layer of ℳ_net_, only the kernels corresponding to the central channels are activated for training. During training, if the validation loss does not decrease after a predefined number of iterations, POI diffusion is triggered. At this point, bisection is recursively applied to determine new active channels, and POI examples are re-merged into their nearest active channels using mean pooling. The kernels corresponding to these newly activated channels then inherit their initial weights from previously activated kernels through nearest copy and are activated for training. This process repeats until all channels are activated.

### Implementation details of DEPAF models

For tasks requiring high localization accuracy and the ability to distinguish overlapping POIs, we upsample both original images and POI examples by a factor of 1.5 using a shared interpolation method, defaulting to cubic spline [16]. In general cases, a factor of 1, i.e., no upsampling, is sufficient. Following this, the training data is normalized to a 0-1 scale using the 99th percentile and the minimum value of the entire training set. It is then randomly sampled into overlapping patches for training. If inhomogeneous background learning is used, the background POI example is set to be three times the size of the central 95% pixel-value distribution area of the signal POI example.

The neural network ℳ_net_ is trained using the Adam optimizer [61] with an initial learning rate of 0.001. The *L*_1_ regularization strength *λ* is warm-started at 1/100 of the target value and increased to the target over 5,000 iterations. Validation loss is measured after every 20 iterations. If the loss does not fall below the historical minimum over 50 consecutive measurements, POI diffusion is triggered and the historical minimum validation loss is reset. If all the channels of the POI examples are activated, the learning rate is halved until it reaches 0.0001. If the loss still does not decrease after another 50 consecutive measurements, the early stopping method [62] is executed, and the model with the lowest historical validation loss is saved as the optimal model.

In the second branch of our method, Bayesian optimization uses the expected improvement (EI) criterion [58, 59] to select the next evaluation point, with a maximum of 80 evaluations. The grid search comprises 100 steps with a step size set to 1/3000 of the pixel value range of *Î*fit. The POI example values in sub-pixel coordinates are obtained using the same interpolation method as in the center-aligned upsampling.

For inferring new images, normalization uses the pre-saved 99th percentile and minimum value from the training set. Since ℳ_net_ uses a fully convolutional architecture, our method can handle images of any size. To prevent memory overflow, images are processed using overlapping sliding windows, with the window size equal to the training patch size and a stride of half the window size. To suppress stitching artifacts caused by edge effects, a Gaussian importance weighting method [63] is used.

Comprehensive testing on multiple datasets demonstrated the high effectiveness of the provided settings, rendering additional user adjustments unnecessary. Users only need to adjust one hyperparameter: the regularization strength *λ* in the loss function for optimizing ℳ_net_. The *λ* determines the model’s noise robustness and POI detection sensitivity. A higher *λ* enhances noise robustness but may miss low-amplitude POIs, and vice versa.

### Acquisition of POI examples

For SMLM, the POI examples are obtained by imaging calibrated fluorescent beads under precisely the same conditions as the sample imaging. In MERFISH analysis, an elliptical Gaussian distribution with different sigma values along the *x* and *y* axes, fitted from the hybridization image, is used as the POI example. For two-photon calcium imaging data denoising, the POI example is determined by averaging the normalized spikes from the original image that fit a single exponential decay function [64]. For active neuron segmentation, a 2D standard Gaussian distribution with a sigma value of 1 is used as the POI example. The POI example used for background estimation benchmarking is a patch of constant value, with a size three times that of the central 95% pixel-value distribution area of the signal POI example.

### Simulating data for performance evaluation

To evaluate the performance of DEPAF, we generate multiple simulated datasets for 2D SMLM benchmarking (Fig. 3a and Extended Data Fig. 2), spike inference benchmarking (Extended Data Fig. 4), denoising evaluation (Extended Data Fig. 5), and background estimation benchmarking (Fig. 6 and Extended Data Fig. 7). This dataset collection spans a wide range of signal densities, SNRs, and background variation scales. Details on the simulation process and parameters can be found in Supplementary Note 2, Supplementary Table 1 and Supplementary Table 2.

### Data processing

#### 2D SMLM benchmarking

Due to computational constraints, DECODE limits the maximum PSF density for simulated data generation in each training epoch. This makes it impractical to use the same high PSF densities as in most of the 18 simulated datasets for training. Therefore, we adopted a zero-shot testing approach to evaluate the performance of DECODE models. Specifically, the models were trained on datasets with a PSF density of 1 PSF/*µ*m^2^ and subsequently tested on datasets with higher PSF densities. This approach reflects a common practical scenario where the PSF density in simulated data may not match the experimental data. For fair comparison, we also conducted zero-shot testing on DEPAF. To ensure there were no systematic errors in the coordinate system definition, such as offsets, *xy*-axis flips, or clear scaling errors, the localization results of each comparison method were manually checked and corrected.

#### 3D SMLM benchmarking

The localization results were directly rendered as scatter plots (Extended Data Fig. 3d) without applying any visual enhancement techniques. The localization results of each comparison method were manually checked and corrected to ensure that there were no systematic errors in the coordinate system definition. To ensure evaluation fairness, we applied the same post-processing as DECODE [4], filtering out low-intensity or high-uncertainty results based on percentage to optimize the results for all methods.

#### Spike inference benchmarking

In spike inference tasks, spike timing is conventionally defined as the onset of the event, i.e., the rising edge of the calcium transient. However, this onset typically does not coincide with the center of the POI, which DEPAF uses as its default localization anchor. To compensate for this systematic offset, we applied a temporal shift to the inferred spike times. For instance, when using a 1×301 spike-shaped POI with the onset positioned at the start of the window, all inferred spike times were shifted 150 samples earlier to align with the true spike onset.

#### Dynamic 2D SMLM

Due to photobleaching, the brightness of fluorescent molecules gradually diminishes over time, hindering the ability of localization algorithms to use consistent sensitivity in detecting single molecules across all SMLM video frames. To address this issue, we employed a histogram-matching-based bleaching correction method [65] to preprocess the collected SMLM videos. This method first analyzes the grayscale histogram of the first frame, then adjusts the grayscale histograms of subsequent frames to match that of the first one, effectively correcting the brightness decay caused by photobleaching and reduces noise increase. Using single-molecule positions obtained from 60 SMLM video frames, we rendered super-resolution frames. To enhance image smoothness, bilinear interpolation was applied when mapping single-molecule positions to pixel values.

#### MERFISH analysis

We applied the histogram-matching-based photobleaching correction method [65] to address brightness fluctuations in MERFISH images across different hybridization rounds and *z*-layers caused by photobleaching. Specifically, using the first *z*-layer of the first hybridization round as a template, we corrected photobleaching across all rounds and *z*-layers, then applied a uniform sensitivity for MERFISH spot signal detection in all images. Compared to the original MERFISH technique, which iteratively adjusted the thresholds (corresponding to sensitivity) in localization algorithms for each round’s images through trial and error [2, 53], this method is more concise.

Each round of hybridization image needs to be registered for barcode decoding. Unlike the original MERFISH technique, which calculated the transformation matrix by fitting and matching the fiducial bead centroids, we found that the entire bead images, including fluorescence background, can be used for registration. Specifically, we first computed the KAZE features points [66] for the bead image corresponding to each round of hybridization image and select the bead image with the most feature points as the template. Then, we calculated the transformation matrix between the feature points of each bead image and those of the template bead image. This method offers greater automation and robustness because the KAZE feature points are more numerous and include descriptors beyond mere positional information.

To accommodate the ultra-high-density MERFISH spot signal detection capability provided by our method, we improved the barcode decoding algorithm to address the more frequent matching conflict issue (i.e., the distance between one spot signal and multiple spot signals is less than the matching threshold). Specifically, we first matched spot signals directly where no conflicts existed and retained all spot signals with matching conflicts. Next, we used hierarchical clustering [67], with the matching threshold as the shortest distance between clusters, to group all conflicting spot signals into clusters. Within each cluster, we applied the Munkres assignment algorithm [68] to obtain the optimal matching that minimizes the overall distance. This method allows ultra-high-density MERFISH spot signal detection results to be decoded into barcodes more accurately while maintaining reasonable time and memory costs and makes the decoding performance less sensitive to the matching threshold.

#### Two-photon calcium imaging

All two-photon calcium imaging videos were aligned non-rigidly using the NoRM-Corre algorithm [69] to eliminate motion artifacts before processing. Each pixel’s time-series values in the video were treated as a 1D image and subjected to median filtering to suppress spike noise. These 1D images were then normalized by dividing them by the difference between their median and first quartile values to address spike amplitude variations between different neurons and cortical regions.

Given the low amount of training data required by our method, we selected and cropped neighborhoods of a few pixels with larger values from these 1D images as potential spike data for training the spike inference model. After training, we obtained noise-free 1D images using our method’s synthesis function and reshaped them back to their original video format, yielding denoised two-photon calcium imaging videos.

Similarly, we selected a few frames with high pixel value sums from the denoised videos as potential active neuron signal data. These frames were used to train the source extraction model based on ultra-high-density PSF localization, using a standard Gaussian PSF as the POI example. After training, to generate the final neuron segmentation masks, we first performed spatial clustering of the localized PSF positions for each frame using the DBSCAN algorithm [56]. For each cluster, we calculated the boundary points, generated the boundary curve using cubic spline interpolation, and extended the boundary curve along its normal direction to compensate for boundary shrinkage caused by the standard Gaussian PSF radius. Finally, we converted each boundary curve into a segmentation mask and merged them using the method employed in SUNS [27], based on criteria such as the mask’s area, center of mass, intersection-over-union (IoU), consume ratio, and the number of consecutive frames.

All parameters, including those for mask generation and merging, were optimized within a pre-set range using Bayesian optimization [58, 59] based on the ground-truth segmentation masks of the training set, aiming to minimize the loss function:

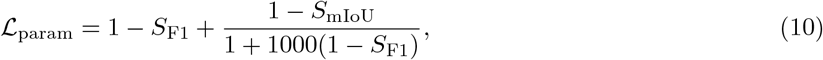

where *S*_F1_ is the F1 score and *S*_mIoU_ is the mean IoU score.

### Evaluation metrics

*SMLM*. The evaluation of SMLM encompasses two aspects: the ability to detect all PSFs and the accuracy of the detected PSF positions. The former is evaluated using the Jaccard index (JI), which is defined as:

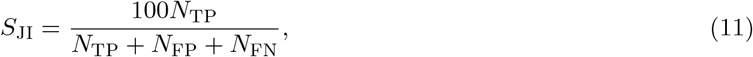

where *N*_TP_ is the true positive rate, *N*_FP_ is the false positive rate, and *N*_FN_ is the false negative rate. The latter is evaluated using the lateral and axial root mean square error (RMSE), which are the distance in nanometers between the localized and the ground-truth positions in lateral and axial direction, respectively. They are defined as:

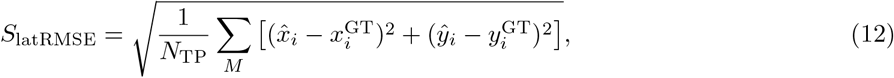

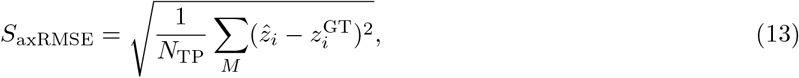

where 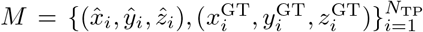 are the matched position pairs within a distance threshold of 1 pixel. The JI and RMSE exhibit a trade-off relationship. To provide a balanced overall assessment of localization performance, the “efficiency” is employed, defined as:

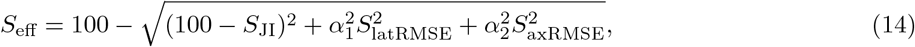

where *α*_1_ and *α*_2_ are weighting parameters set to 1 nm^−1^ and 0.5 nm^−1^, respectively [36].

#### Spike inference

The evaluation of spike inference is performed using two metrics that characterize the timing accuracy and overlap between the ground-truth and inferred spike trains: Victor–Purpura distance [40] and error rate. Let *A* ={ *a*_1_, *a*_2_, …, *a*_*m*_} and *B* = {*b*_1_, *b*_2_, …, *b*_*n*_} denote the ground-truth and inferred spike trains, respectively. The Victor–Purpura distance quantifies the minimum cost of transforming one spike train into another using spike insertion (cost 1), deletion (cost 1), or temporal shifting (cost equals the time difference in seconds). It is computed via dynamic programming and defined as:

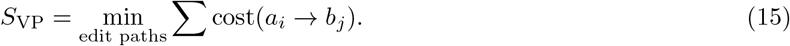

To assess set-level matching, we define a match as an inferred spike time that occurs within a tolerance of 0.5 s from a ground-truth spike time. Based on the number of matched spikes *N*_match_, the error rate is calculated as:

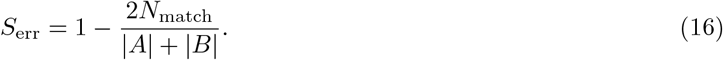

#### Background estimation

The evaluation of background estimation employs three metrics to assess both the accuracy of pixel-wise background reconstruction and the fidelity of the foreground signals following background removal: background RMSE, background correlation, and peak signal-to-noise ratio (PSNR) of background-removed signal images. Prior to evaluation, all background and background-subtracted images are min-max normalized to the range [0, 1] on a per-frame basis. The background RMSE is defined as:

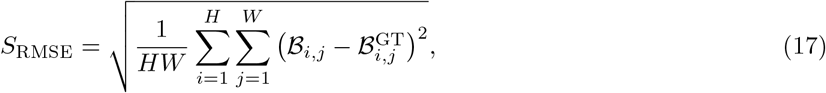

where *H* and *W* denote the height and width of the image, and ℬ and ℬ^GT^ denote the estimated and ground-truth background images, respectively. The background correlation is quantified as:

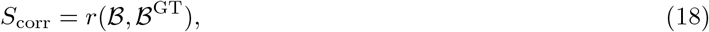

where *r*(·, ·) denotes the Pearson correlation coefficient function. The PSNR is defined as:

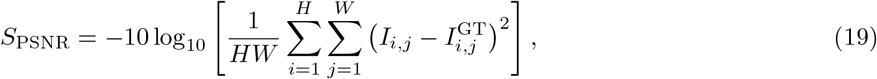

where *I* and *I*^GT^ represent the background-removed and ground-truth signal image, respectively.

#### MERFISH analysis

Evaluating MERFISH analysis involves assessing the accuracy of RNA expression quantification and barcode decoding. The accuracy of RNA expression quantification is determined by the Pearson correlation coefficient between the logarithmic scale of the copy number of each RNA species detected through MERFISH and the abundance determined by FPKM. It is defined as:

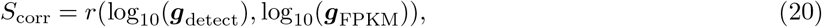

where ***g***_detect_ is the vector of the copy number of each RNA species detected through MERFISH, ***g***_FPKM_ is the vector of the abundance of each RNA species determined by bulk sequencing in FPKM, and *r*( ·, ·) is the Pearson correlation coefficient function. Since the codebook uses MHD4 code [2], a match is determined if the Hamming distance between the decoded barcode and any barcode in the codebook is less than or equal to 1. The accuracy of barcode decoding is evaluated by the average per-bit 0 *→* 1 and 1 *→* 0 error rates in the matched barcodes. The average per-bit 0 *→* 1 error rate is defined as:

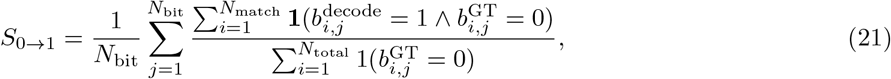

and the average per-bit 1 *→* 0 error rate is defined as:

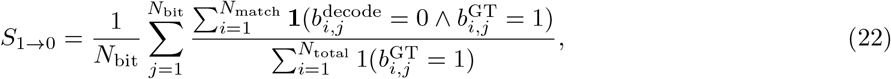

where *N*_match_ is the total number of matched barcodes, *N*_bit_ is the number of bits in the codebook, *b*^decode^ is the *j*-th bit of the *i*-th decoded barcode, *b*^GT^ is the *j*-th bit of the ground-truth barcode matched with the *i*-th decoded barcode, and **1**(·) is the indicator function, which returns 1 if the input is true and 0 otherwise.

#### Active neuron segmentation

The neuron segmentation results are evaluated by comparing them with ground-truth labels. To match the segmentation masks with the ground-truth labels, the distance between a ground-truth mask *M* ^GT^ and a detected mask *M*_*j*_ is determined based on their IoU score as follows:

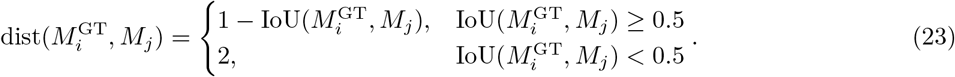

Here,

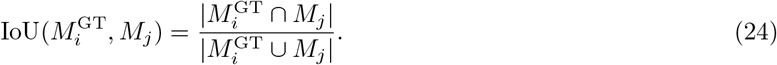

In this formula, a distance of 2 is assigned to mask pairs with an IoU score below 0.5, indicating that these masks are too dissimilar to be considered matches. The Munkres assignment algorithm [68] is then applied to this distance matrix to solve the linear assignment problem, identifying matched mask pairs where the distance is less than 2 as true positives. Next, the performance is quantified using F1 score defined as:

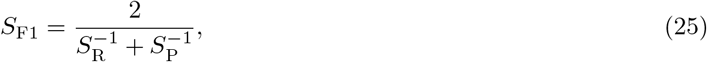

and the mean IoU score defined as:

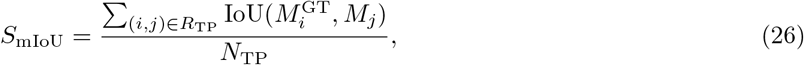

where *S*_R_ = *N*_TP_*/N*_GT_ denotes recall, *S*_P_ = *N*_TP_*/N*_detect_ denotes precision, *R*_TP_ is the set of true positive pairs, *N*_TP_ is the number of true positives, *N*_GT_ is the total number of ground-truth labels, and *N*_detect_ is the number of neurons detected by the evaluated methods.

### Sample preparation and imaging of dynamic 2D SMLM

See Supplementary Note 3 for details on the sample preparation and imaging of dynamic 2D SMLM.

## Benchmarking algorithms

For 2D SMLM benchmarking, we used DECODE [4] (v0.10.2, https://github.com/TuragaLab/DECODE), SMAP [15] (v201217, https://github.com/jries/SMAP), CSpline [16] (v2.2, https://github.com/ZhuangLab/storm-analysis), rainSTORM [17] (v3.1.8, https://titan.physx.u-szeged.hu/ ∼adoptim/?page id=582), and rapidSTORM [18] (v3.3.1, https://github.com/stevewolter/rapidstorm). For dynamic 2D SMLM, we again used DECODE and SMAP, and additionally included DBlink [5] (https://github.com/alonsaguy/Dblink) and eSRRF [37] (https://github.com/HenriquesLab/NanoJ-eSRRF). For 3D SMLM benchmarking, we again used DECODE, SMAP, and CSpline, and further included QC-STORM [20] (v3.7.5.2, https://github.com/SRMLabHUST/QC-STORM) and Easy-DHPSF [19] (v2.1, https://sourceforge.net/projects/easy-dhpsf). For MERFISH analysis, we used HSCMERFISH [54] (https://campuspress.yale.edu/wanglab/HSCMERFISH/). For spike inference benchmarking, we used MLspike [22] (https://github.com/MLspike/spikes), Peeling [23] (v0.1, https://github.com/HelmchenLab/CalciumSim), FastL0 [24] (https://github.com/jewellsean/FastLZeroSpikeInference), OASIS [21] (https://github.com/zhoupc/OASIS), and CASCADE [29] (v2.0, https://github.com/HelmchenLabSoftware/Cascade). For background estimation benchmarking, we used Dark Sectioning [70] (v1.0, https://github.com/Cao-ruijie/Dark-sectioning), Sparse Deconvolution [71] (v1.0.3, https://github.com/WeisongZhao/Sparse-SIM), WBNS [72] (https://github.com/NienhausLabKIT/HuepfelM), and the Fiji plugins Sliding Paraboloid [73] and Rolling Ball [74]. For active neuron segmentation, we used CaImAn batch [25] (v1.6.4, https://github.com/flatironinstitute/ CaImAn), Suite2p [26] (v0.6.16, https://github.com/cortex-lab/Suite2P), SUNS batch [27] (v1.1.1, https://github.com/YijunBao/Shallow-UNet-Neuron-SegmentationSUNS), and STNeuroNet [28] (https://github.com/soltanianzadeh/STNeuroNet). Their parameter settings are provided in Supplementary Note 4.

## Data available

The data and trained models for 2D SMLM localization benchmarking (Fig. 3a and Extended Data Fig. 2), dynamic 2D SMLM (Fig. 3b,c and Fig. 6d), spike inference benchmarking (Extended Data Fig. 4), denoising evaluation (Extended Data Fig. 5) and background estimation benchmarking (Fig. 6a,b and Extended Data Fig. 7) are available at https://zenodo.org/records/15862305. The full-length supplementary video 2 are available at https://zenodo.org/records/14145974. The 3D SMLM benchmarking dataset MT0.N2.HD-DH (Extended Data Fig. 3) is publicly available at https://srm.epfl.ch/srm/dataset/challenge-3D-simulation/ MT0.N2.HD/index.html. The video J123 (Fig. 5 and Extended Data Fig. 6) is publicly available at https://zenodo.org/records/1659149. Raw data for MERFISH analysis (Fig. 4 and and Fig. 6c) are available upon request from the authors of ref. [54]. All other data supporting this study are available from the corresponding authors upon reasonable request.

## Code available

DEPAF is accessible as Supplementary Software. The latest version can be found at https://github.com/zhang-fengdi/DEPAF. The code to reproduce the results in this paper can be accessed at https://github.com/zhang-fengdi/DEPAF/tree/main/paper_reproduction.

## Acknowledgements

J.G. was supported by the National Key Research and Development Program of China (Grant No. 2022YFA1103401). R.H. was supported by the National Key Project (Grant No. 62331006). Y.F. was supported by the National Key R&D Program of China (Grant No. 2022YFC3300704) and the National Key Project (Grant No. 62331006). We thank Prof. Minping Qian from Peking University for her helpful discussions and suggestions.

## Author contributions

R.H. and J.G. initiated the project. F.Z. developed the software, analyzed the data, and drafted the manuscript with contributions from all authors. R.H., J.G., H.M and D.G. contributed to result interpretation. J.G. and M.X. provided data support. Y.F. assisted with analysis and result discussions. R.H., J.G. and X.J. supervised the project and the final manuscript.

## Competing interests

F.Z., J.G., R.H. and M.X. have a pending patent application on the presented method. The other authors declare no competing interests.

**Extended Data Fig. 1.**
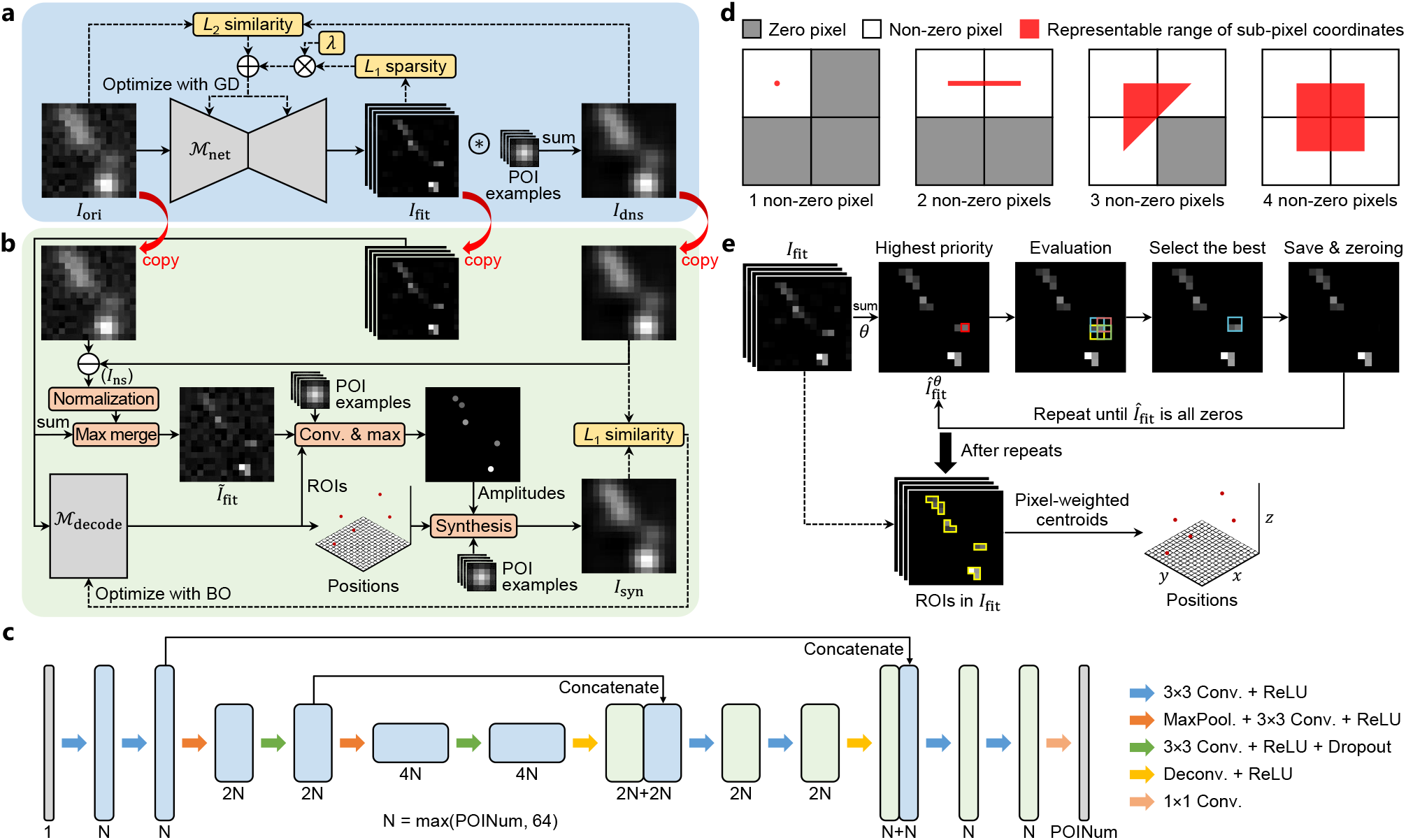
Method detailed overview. **a**, The first branch of our method. GD, gradient descent. **b**, The second branch of our method. Conv., convolution; Max, maximum; BO, Bayesian optimization. **c**, Architecture of ℳ_net_ with an encoder depth of 2 as an example. This neural network comprises a down-sampling stage followed by an up-sampling stage. Each stage consists of several (de)convolutional layers with 3×3 filters, along with non-trainable operations including max-pooling (stride of 2), dropout (probability of 50%), and rectified linear unit (ReLU) activation. The first convolutional layer in the down-sampling stage has *N* filters, where *N* is the maximum of 64 and the number of POI examples. In each down-sampling phase, the resolution is halved while the number of filters is doubled; the reverse is true for each up-sampling phase. Black arrows denote skip connections. POINum, the number of POI examples; Conv., convolutional layer; MaxPool., max-pooling layer; Deconv., deconvolutional layer. **d**, Schematic depiction of the priors used in ℳ_decode_. For ROIs containing 1 to 4 non-zero pixels, their representable range of sub-pixel coordinates in continuous coordinate space expands progressively. According to classical probability theory, their occurrence probability increases correspondingly, resulting in ascending partitioning priority ordered by the number of non-zero pixels. **e**, Schematic depiction of the processing procedure for ℳ_decode_. Starting from the image 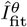 obtained by summing the channels of *I*_fit_ and filtering it with threshold *θ*, the algorithm iteratively selects pixels with the minimal number of neighbors, computes pixel sums in surrounding 2×2 blocks, selects and saves the block with the largest pixel sum, and updates the image until all ROIs are partitioned.

**Extended Data Fig. 2.**
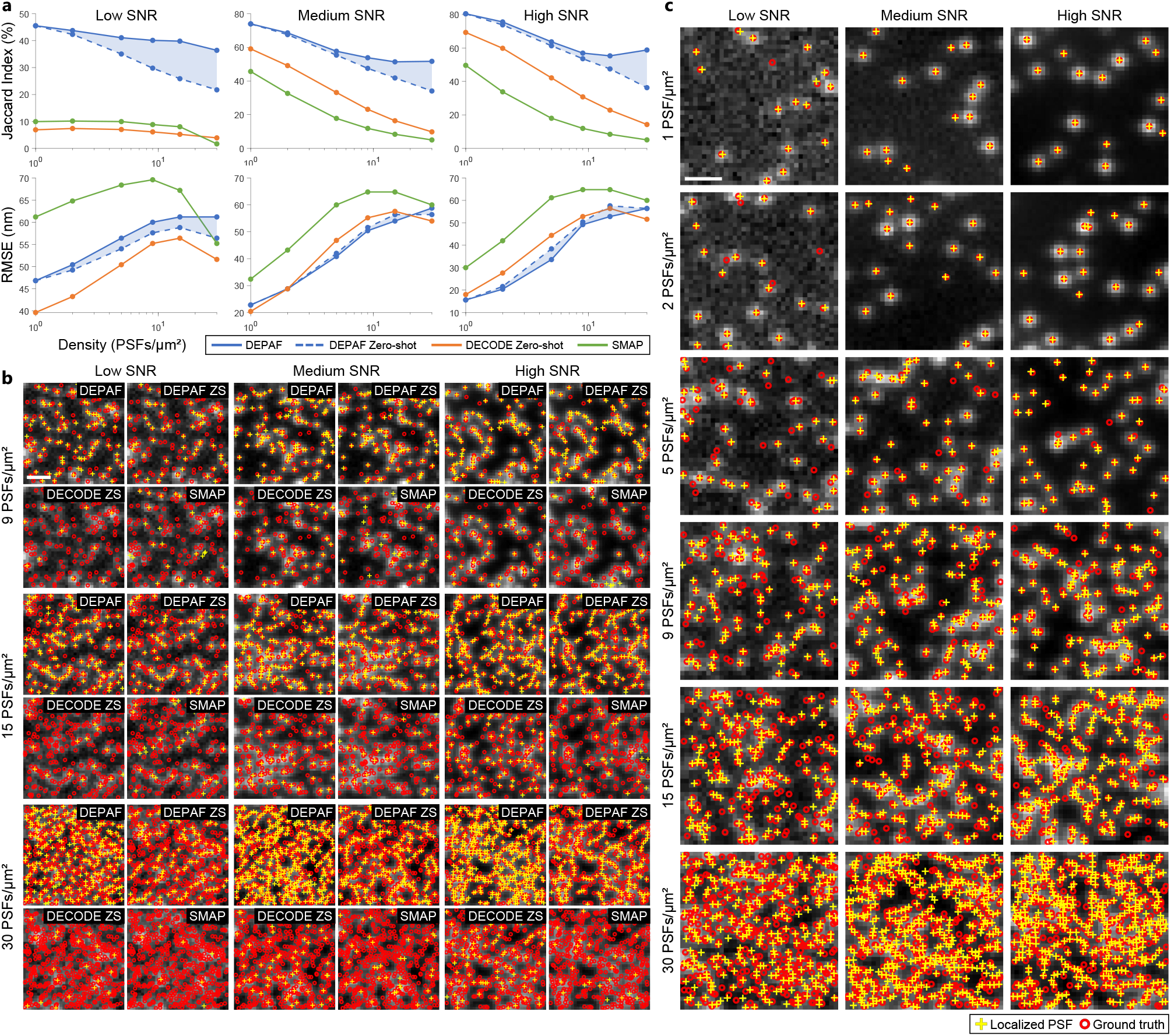
Detailed comparison results of localization performance across different methods on 18 simulated SMLM datasets. **a**, The line charts compares the Jaccard index and RMSE of different methods across different SNR levels and PSF densities (log scale) in 18 simulated datasets. **b**, Each column of the images visualizes the localization results of various methods at PSF densities of 9, 15, and 30 PSFs/*µ*m^2^ for low, medium, and high SNR levels, respectively. Scale bar, 1 *µ*m. ZS, zero-shot testing. **c**, Each column of images provides a more detailed visualization of DEPAF’s localization results at PSF densities of 1, 2, 5, 9, 15, and 30 PSFs/*µ*m^2^, corresponding to low, medium, and high SNR levels. Scale bar, 1 *µ*m.

**Extended Data Fig. 3.**
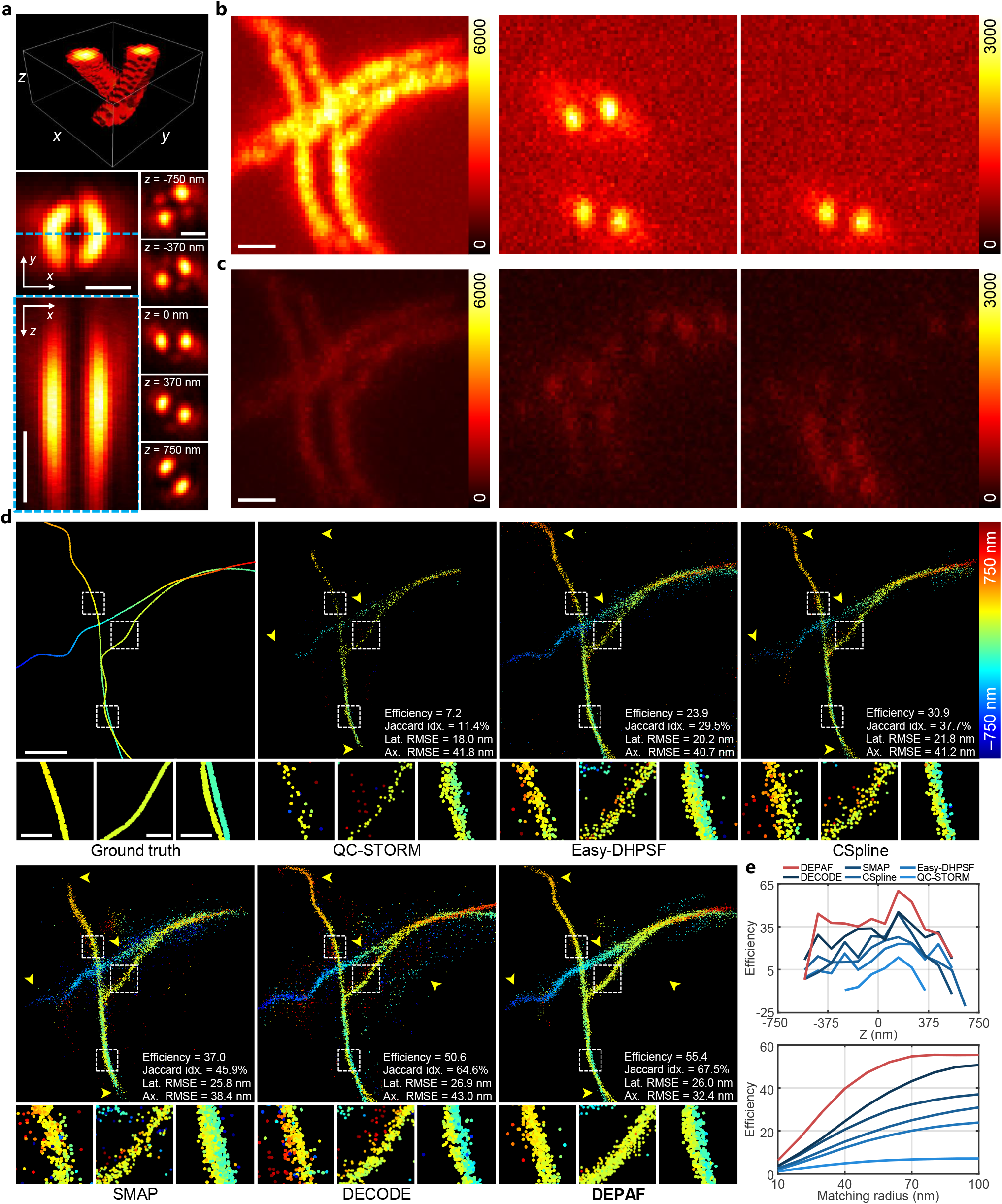
DEPAF enables high-robust-accuracy 3D SMLM in imaging conditions with low SNR and high PSF density. **a**, 3D view of real double-helix PSFs acquired using calibrated beads at 151 different *z*-axis positions (ranging from −750 nm to 750 nm, with a 10 nm interval) for generating the simulated dataset (top). Maximum intensity projection of the double-helix PSFs in the *xy* view (middle left). Scale bar, 1 *µ*m. Intensity profile in the *xz* view along the blue dashed line in the maximum intensity projection (bottom left). Scale bar, 500 nm. Representative PSF images at different *z*-axis positions (bottom right). Scale bar, 1 *µ*m. **b**, Temporal maximum intensity projection (left) and example frames (middle and right) from 2,500 simulated frames showing typical Alexa647-labeled STORM data [36]. Scale bar, 1 *µ*m. **c**, Data used for reconstruction, with 10 times the PSF density and an average photon count reduced to 10.8% of that in **b**. Scale bar, 1 *µ*m. **d**, Ground-truth 3D super-resolution image for generating the simulated dataset, and the reconstructed image using QC-STORM, Easy-DHPSF, CSpline, SMAP, DECODE and DEPAF. Yellow arrows indicate specific regions to highlight differences in resolution and structural details among the methods. Metrics, computed with a 100 nm matching radius, are shown in the bottom right of each image. Scale bar, 1 *µ*m. Idx., index. Lat., lateral. Ax., axial. Zoomed-in views of the boxed regions present detailed visualization of the reconstructions. Scale bar, 200 nm. **e**, Top: Efficiency across *z* -slices with bin size 100 nm, evaluated at a 100 nm matching radius. Bottom: Efficiency as a function of matching radius, showing performance at different evaluation tolerances.

**Extended Data Fig. 4.**
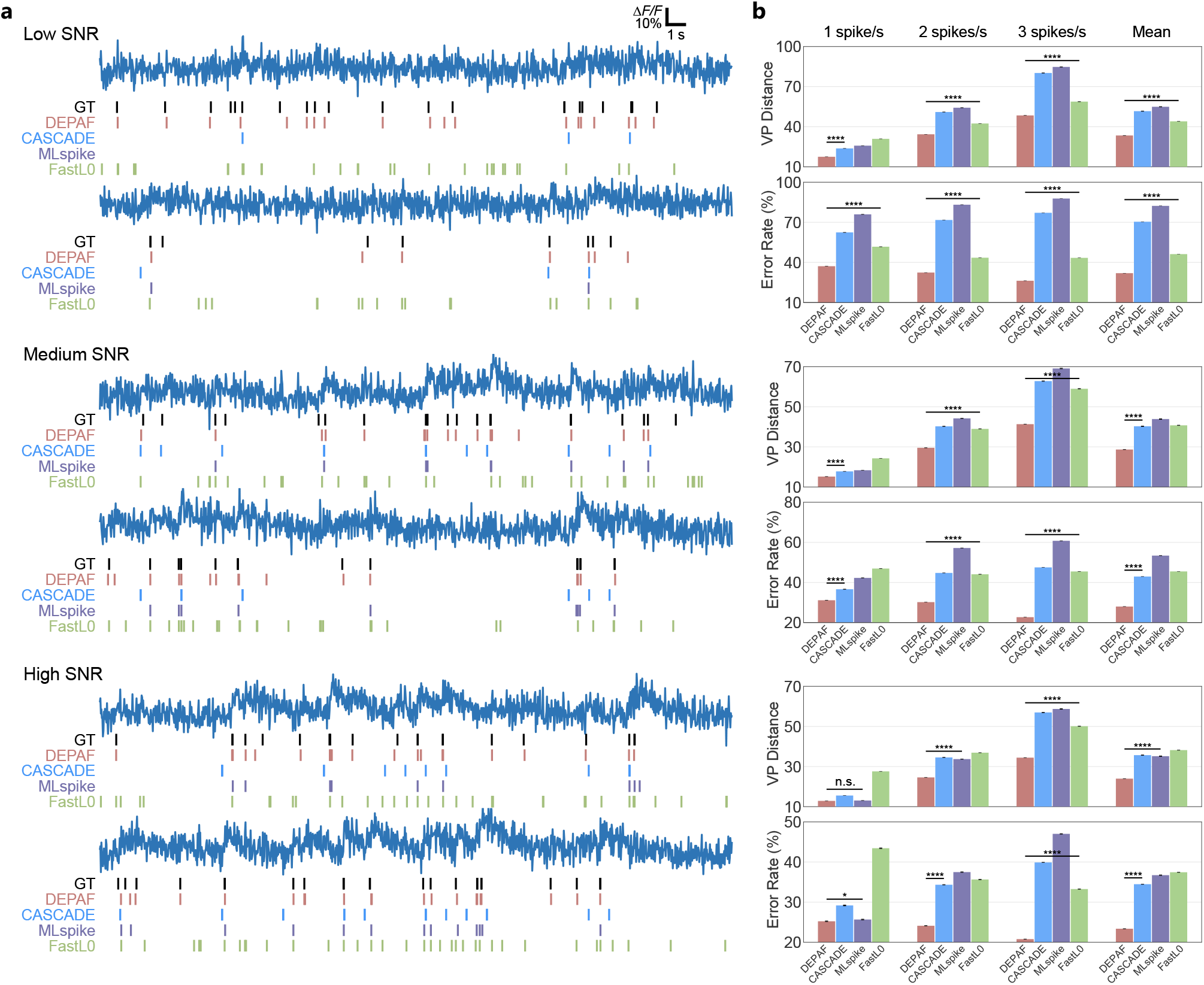
DEPAF outperforms other methods in spike inference, particularly under low SNR conditions. **a**, Performance of different spike inference methods, including DEPAF, CASCADE, MLspike, and FastL0, on data with low (first and second rows), medium (third and fourth rows), and high (fifth and sixth rows) SNR levels at a spike rate of 1 spike/s. Vertical lines represent spike start times. Δ*F* represents the change in fluorescence relative to the baseline fluorescence *F*_0_, which is approximated by the average of the entire trace. **b**, Performance quantification at low, medium, and high SNR levels using VP distance and error rate for spike rates of 1, 2, and 3 spikes/s, and their average. *: *P <* 0.05, ****: *P <* 0.0001, n.s.: not significant. *P* values are calculated using the two-sided Wilcoxon signed-rank test. Error bars represent SEM (*n* = 2,000 traces).

**Extended Data Fig. 5.**
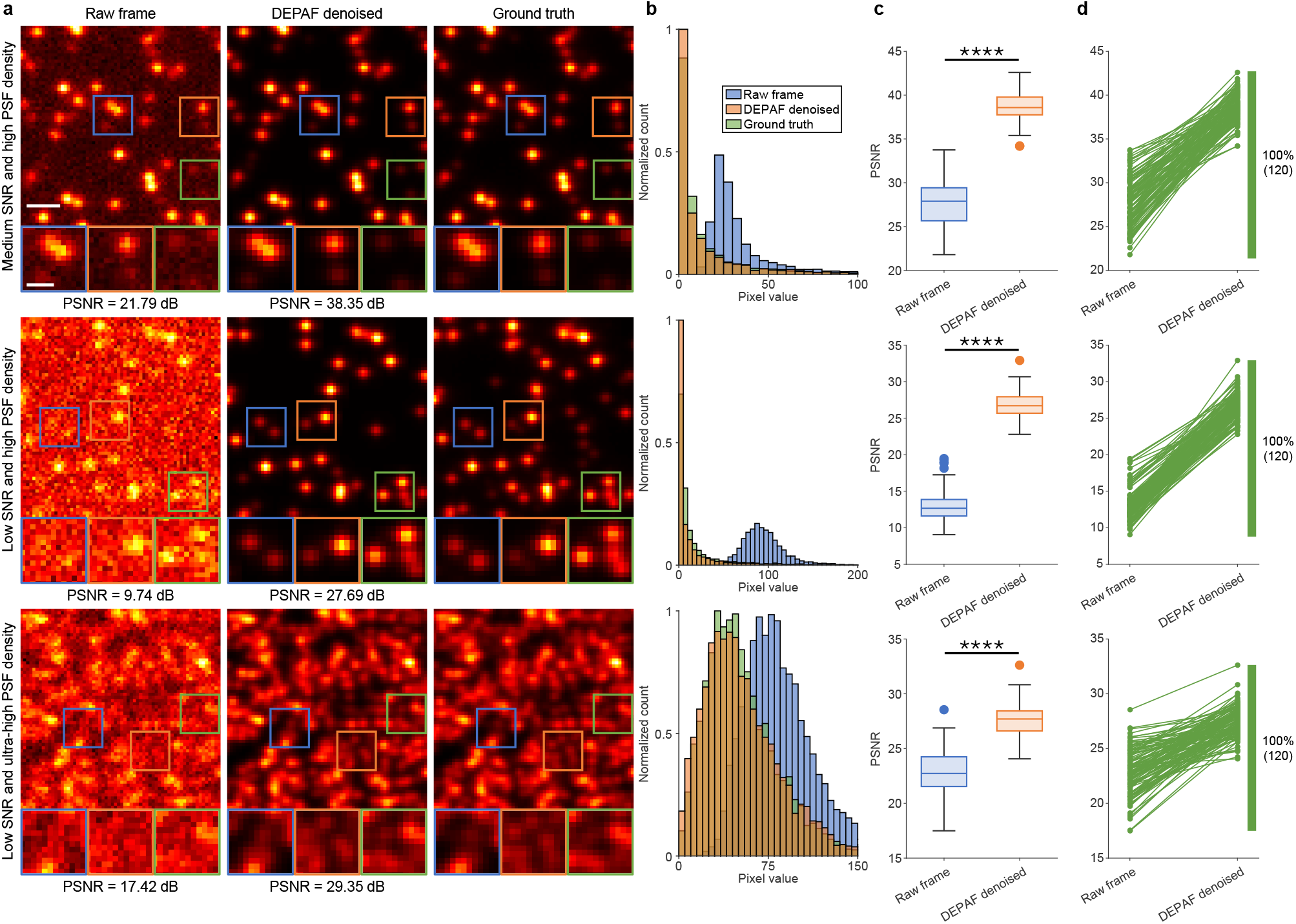
DEPAF achieves strong denoising performance across different SNRs and PSF densities through its image synthesis function, benefiting from its accurate estimation of POI positions and amplitudes. **a**, Comparison of simulated data in raw frames, DEPAF denoised frames, and ground truth. Each row shows images under different SNR and PSF densities. Scale bar, 1 *µ*m. The subplots below display magnified details of the denoising results. Scale bar, 500 nm. The peak signal-to-noise ratio (PSNR) is indicated below each raw frame and DEPAF denoised frame. **b**, Histograms of pixel values under different conditions, comparing the distributions of input frames (blue), denoised frames (orange), and ground truth (green). **c**, Box plots of PSNR values for each condition, showing the performance difference between original images and DEPAF denoised images. *P* values calculated by one-sided paired *t*-test are specified with asterisks; \*\*\*\**P <* 0.0001 for all comparisons. **d**, Comparison of PSNR between original images and denoised images under different conditions, with PSNR markedly improved in each condition.

**Extended Data Fig. 6.**
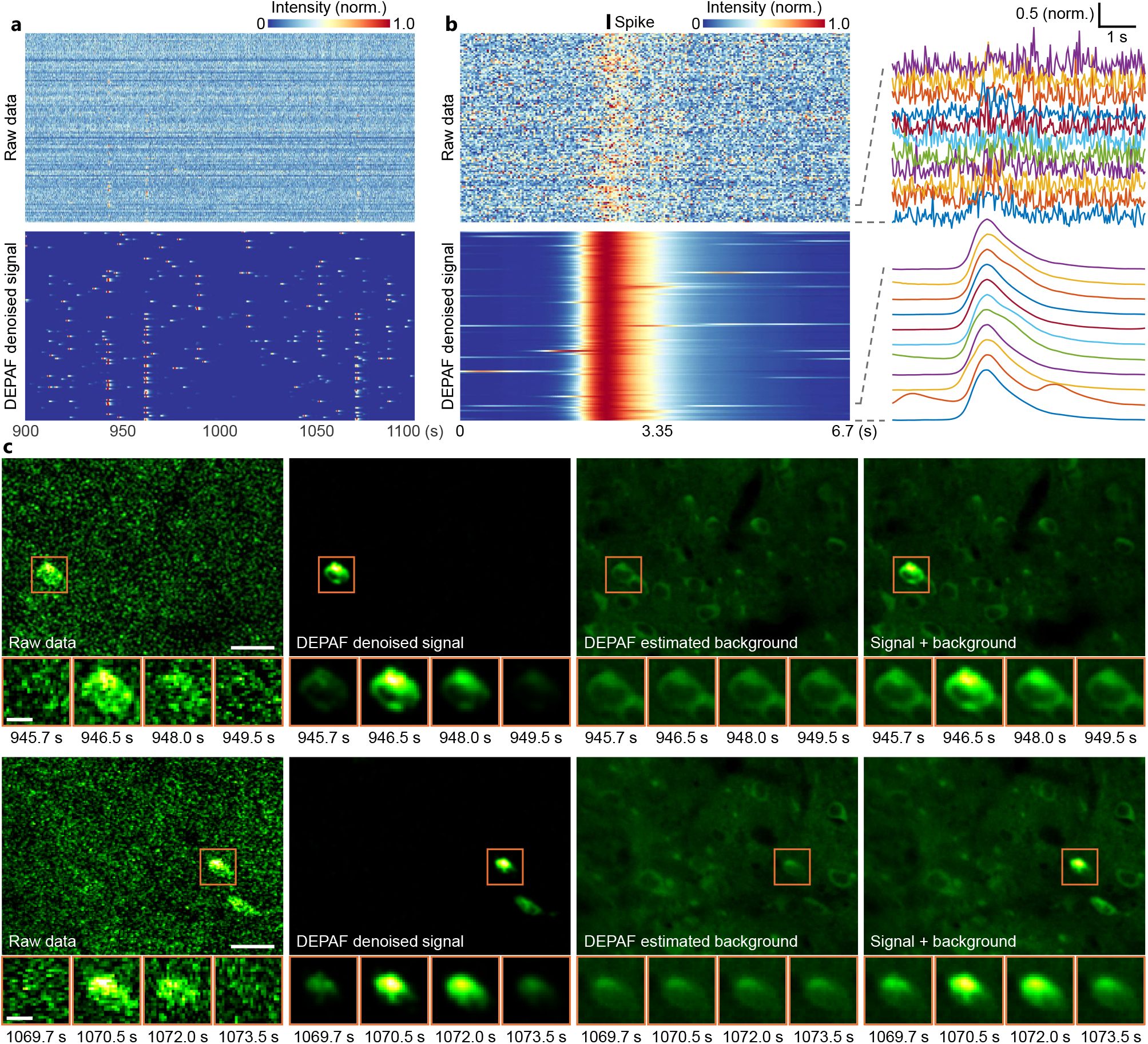
Extended results of DEPAF in denoising and background separation for two-photon calcium imaging videos. **a**, Fluorescence traces from 120 pixels. Top, raw recording. Bottom, background-separated signals denoised by DEPAF. Norm., normalized. **b**, Calcium fluctuations induced by 120 action potentials. All spikes were normalized and temporally aligned to the black bar. Zoomed-in traces are displayed in the right panel. Norm., normalized. **c**, First column: raw somatic signal recording. Second column: background-separated signals denoised by DEPAF. Images were spatially smoothed using a standard Gaussian kernel. Third column: backgrounds estimated by DEPAF. Fourth column: sum of backgrounds and signals. Scale bar, 20 *µ*m. Zoomed-in views of the boxed regions present calcium transients within a 3.8-second time window. Scale bar, 6 *µ*m.

**Extended Data Fig. 7.**
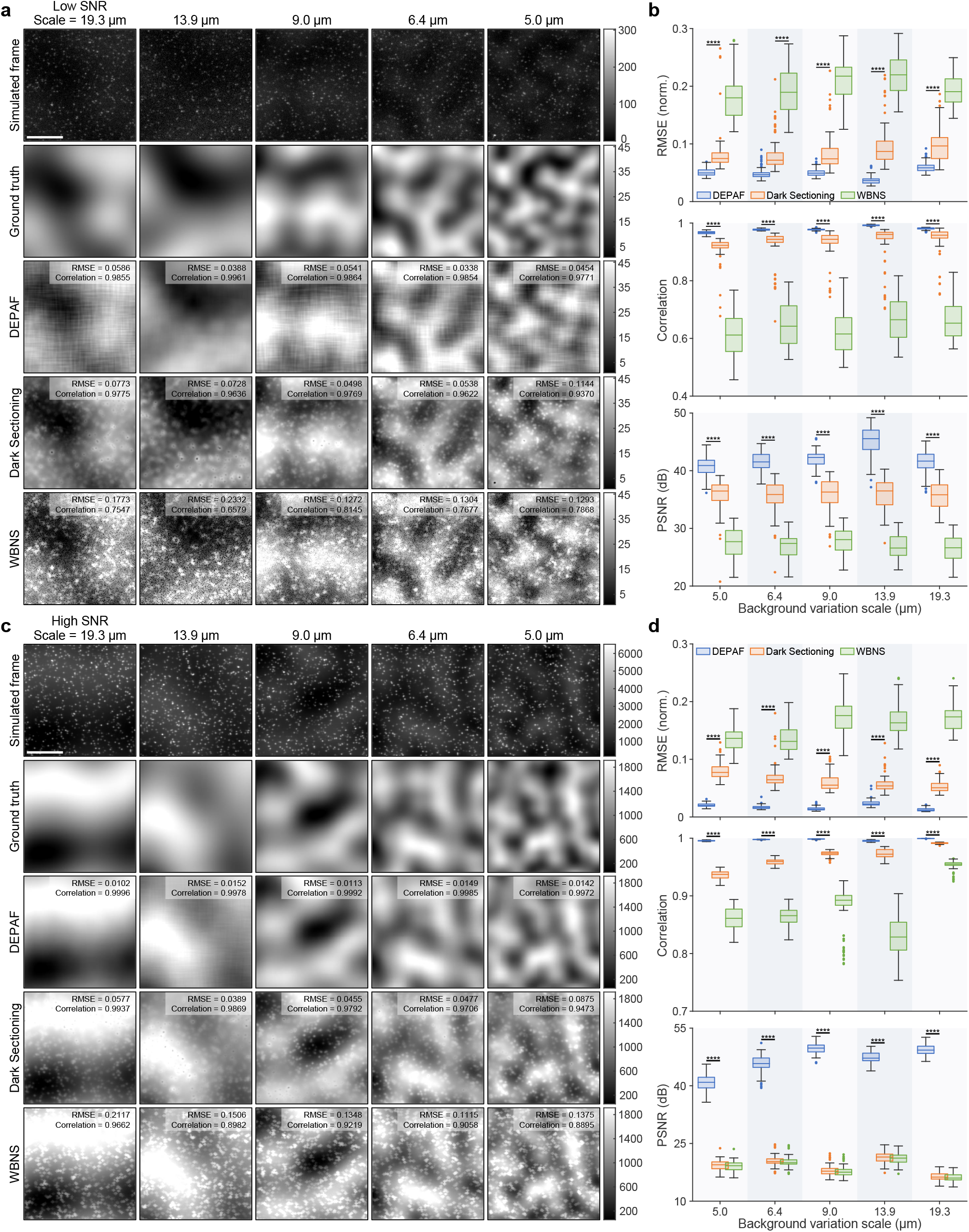
Extended results for DEPAF in inhomogeneous background estimation under low and high SNR levels. **a**, Performance comparison of DEPAF, Dark Sectioning, and WBNS under low SNR conditions and varying background variation scales. First row: simulated fluorescence frames. Second row: ground-truth backgrounds. Rows three to five: backgrounds estimated by DEPAF, Dark Sectioning, and WBNS, respectively. RMSE and Pearson correlation coefficients with ground truth are shown in the top right of each image. Scale bar, 10 *µ*m. **b**, Quantitative comparison of RMSE, Pearson correlation coefficients, and PSNR of background-removed images under low SNR across background variation scales. \*\*\*\**P <* 0.0001 for all comparisons, determined using two-sided *t*-tests. Norm., normalized. **c**,**d**, Same analyses as in **a** and **b**, respectively, but under high SNR conditions.

