## Supplementary Information for "Self-contrastive learning enables interference-resilient and generalizable fluorescence microscopy signal detection without interference modeling"

[2]Peng Cheng Laboratory, Shenzhen, 518055, China.

**5 Supplementary tables** **8**

**6 Supplementary figures** **10**

**7 Supplementary video descriptions** **15**

**References** **18**

### 1 Implementation of center-aligned upsampling

Considering a POI example with a height $H$ and width $W$, its coordinate vector of the pixels along the $x$-axis is:

$$\boldsymbol{x}_{\mathrm{ori}} = (0, 1, \ldots, W-1). \tag{1}$$

Let the initial upsampling factor be $k > 1$. To ensure that the edge lengths of the POI example remain odd after upsampling, which ensures a well-defined center for convolution operations, we first correct the upsampling factor:

$$\hat{k} = \begin{cases} \frac{\lceil kW \rceil + 0.5}{W}, & \text{if } \lceil kW \rceil \bmod 2 = 0 \\ k, & \text{otherwise} \end{cases}. \tag{2}$$

Next, we expand the $\boldsymbol{x}_{\mathrm{ori}}$ to $\boldsymbol{x}_{\mathrm{exp}}$ with an interpolation step size $s = 1/\hat{k}$ and an upsampled width $\lceil \hat{k}W \rceil$ by:

$$\boldsymbol{x}_{\mathrm{exp}} = (0, s, 2s, 3s, \ldots, \lceil \hat{k}W \rceil s - s). \tag{3}$$

We then perform center alignment on $\boldsymbol{x}_{\mathrm{ori}}$ and $\boldsymbol{x}_{\mathrm{exp}}$ to obtain the $x$-axis coordinates $\boldsymbol{x}_{\mathrm{qry}}$ of the interpolation query points by:

$$\boldsymbol{x}_{\mathrm{qry}} = \boldsymbol{x}_{\mathrm{exp}} + (\bar{x}_{\mathrm{ori}} - \bar{x}_{\mathrm{exp}}), \tag{4}$$

where $\bar{x}_{\mathrm{ori}} = (W-1)/2$ and $\bar{x}_{\mathrm{exp}} = (\lceil \hat{k}W \rceil s - s)/2$ are the mean values of $\boldsymbol{x}_{\mathrm{ori}}$ and $\boldsymbol{x}_{\mathrm{exp}}$, respectively. Similarly, the $y$-axis coordinates $\boldsymbol{y}_{\mathrm{qry}}$ of interpolation query points can be calculated through the same procedure. Then the coordinates of interpolation query points are structured by a full grid of $\boldsymbol{x}_{\mathrm{qry}}$ and $\boldsymbol{y}_{\mathrm{qry}}$ as:

$$Q = \begin{bmatrix} (x^1_{\mathrm{qry}}, y^1_{\mathrm{qry}}) & \cdots & (x^{\lceil \hat{k}W \rceil}_{\mathrm{qry}}, y^1_{\mathrm{qry}}) \\ \vdots & \ddots & \vdots \\ (x^1_{\mathrm{qry}}, y^{\lceil \hat{k}H \rceil}_{\mathrm{qry}}) & \cdots & (x^{\lceil \hat{k}W \rceil}_{\mathrm{qry}}, y^{\lceil \hat{k}H \rceil}_{\mathrm{qry}}) \end{bmatrix}. \tag{5}$$

The upsampling process for the original images follows the same method as that applied to the POI examples with the corrected upsampling factor $\hat{k}$. Thus, after performing POI localization on the upsampled images, the POI positions can be mapped back to the original coordinate system by:

$$\hat{x}_{\mathrm{ori}} = (\hat{x}_{\mathrm{up}} - \lceil \hat{k}W \rceil/2)/\hat{k} + \left\lfloor \lceil \hat{k}W \rceil/\hat{k} \right\rfloor/2, \tag{6}$$

where $\hat{x}_{\mathrm{up}}$ is the $x$-axis coordinate of a POI in the upsampled image. Similarly, $\hat{y}_{\mathrm{ori}}$ can be calculated accordingly.

### 2 Test data simulation

#### 2.1 2D SMLM benchmarking

We used the data simulation module of DECODE [1] to generate 18 simulated datasets with high or ultra-high PSF densities of 1, 2, 5, 9, 15, and 30 PSFs/$\mu$m$^2$ under three different SNR levels: high (average PSF intensity of 40,000 with a standard deviation of 8,000), medium (7,000 $\pm$ 3,000), and low (1,000 $\pm$ 400). All other settings were kept fixed (Supplementary Table 1). To better simulate real imaging conditions, we added inhomogeneous backgrounds, which were generated using Perlin noise [2], with an average variation scale of 12.5 $\mu$m and maximum intensities at $\sim$30% of the corresponding average PSF intensities. The background variation scale was determined by the distance between the nearest peak and trough in the backgrounds.

#### 2.2 Spike inference benchmarking

We used the simulation tools provided by MLspike [3] to generate 9 simulated datasets of neuronal calcium fluorescence traces across three spike rate conditions (1, 2, and 3 Hz) and three SNR levels: high ($\Delta$F/F amplitude of 0.07–0.075), medium (0.05–0.055), and low (0.03–0.035). Each dataset consists of 2,000 traces. The calcium signal dynamics were defined by a decay time constant $\tau$ and noise level $\sigma$, randomly sampled from log-uniform distributions within the ranges of [0.999, 1.000] and [0.045, 0.050], respectively. To better simulate realistic imaging conditions, we incorporated baseline drift into the traces. All other simulation parameters were kept fixed (Supplementary Table 2).

#### 2.3 Background estimation benchmarking

The inhomogeneous backgrounds were generated using Perlin noise [2] with average variation scales of 5.0, 6.4, 9.0, 13.9, and 19.2 $\mu$m, defined as the distance between the nearest peak and trough. These backgrounds were then added to the simulated data generated by the DECODE data simulation module with a PSF density of 1 PSF/$\mu$m$^2$. The PSF intensities for the datasets with "low, medium, and high SNR" were 1,000 $\pm$ 400, 7,000 $\pm$ 3,000, and 40,000 $\pm$ 8,000, respectively. All other settings were kept fixed (Supplementary Table 1). The maximum intensities of the added backgrounds are $\sim$30% of the corresponding average PSF intensities.

#### 2.4 Denoising performance evaluation

The simulated SMLM dataset for evaluating denoising performance was generated using the data simulation module of DECODE. In this dataset, the PSF density is 1 PSF/$\mu$m$^2$ for "high PSF density" data and 15 PSFs/$\mu$m$^2$ for "ultra-high PSF density" data. The PSF intensities for the low-SNR and medium-SNR raw data are 1,000 $\pm$ 400 and 7,000 $\pm$ 3,000, respectively, while the corresponding ground-truth data have average intensities 100 times and 57 times higher. The PSF positions in the ground-truth data are set to be the same as those in the raw data. All other settings were kept fixed (Supplementary Table 1).

### 3 Sample preparation and imaging of dynamic 2D SMLM

COS-7 cells (ATCC, CRL-1651) were cultured in Dulbecco's modified eagle medium (DMEM) (Invitrogen, #11965-118) supplemented with 10% fetal bovine serum (Gibco, #16010-159) and 100 $\mu$g/mL penicillin and streptomycin (Invitrogen, #15140122) to prevent bacterial contamination. The cells were maintained under standard conditions of 5% $CO_2$, a humidified environment, and 37°C. To optimize cell culture conditions, glutamine and non-essential amino acids were added.

During cell passaging, the cells were washed three times with pre-warmed phosphate-buffered saline (PBS) (Life Technologies, #14190500BT) and then digested with 0.25% trypsin (Gibco, #25200-056) for 30 s. Cells were seeded at a density of $5 \times 10^4$–$10 \times 10^4$ cells/mL in 35 mm, #1.5 glass-bottom culture dishes (SunBloss™, STGBD-035-1). The cells were tested for mycoplasma contamination using MycoAlert (Lonza), and all tests returned negative results.

After 48 hours of stable growth, transfection was performed. On the day of transfection, 7.5 $\mu$L of Lipofectamine™ 3000 reagent (Invitrogen, #L3000001) was diluted in 125 $\mu$L of Opti-MEM™ medium (Invitrogen, #31985070) and mixed well. Separately, 5 $\mu$g of MAP4-HaloTag fusion DNA was diluted in 115 $\mu$L of Opti-MEM™ medium, followed by the addition of 10 $\mu$L of P3000™ reagent, and mixed evenly. The diluted DNA and Lipofectamine™ 3000 reagent were combined at a 1:1 ratio and incubated for 15 min to form DNA-lipid complexes. These complexes were then added to the cells, which were cultured in serum-free medium for 48 to 56 hours. Subsequently, 200 nM Janelia Fluor® HaloTag® Ligand JF646 (GA1120) was added to the cells and incubated for 30 min at 37°C in a 5% $CO_2$ incubator. After incubation, the medium was aspirated and replaced with fresh medium. Finally, the cells were transferred to a microscope for imaging using Hank's balanced salt solution (HBSS) containing 200 nM Trolox as a live-cell imaging buffer.

Imaging experiments were conducted using an Abbelight SAFe 180 system (Abbelight, France) mounted on an Olympus IX83 microscope, equipped with 532-nm and 640-nm Oxxius lasers and a Hamamatsu Orca Fusion v3 sCMOS camera, using a 100× oil-immersion objective (NA 1.49). Imaging parameters were set to a camera sensor resolution of 256×256 pixels, pixel size of 108 nm, and imaging speed of 1.28 ms/frame. Imaging was performed using the 640 nm laser at 5% power.

### 4 Parameter settings for benchmarking algorithms

To ensure fair and consistent comparisons, we identified which parameters could remain at default values and which required tuning, based on official documentation and empirical testing. Tunable parameters were optimized using a coarse-to-fine grid search. Below, we list the parameter settings used for each method across the evaluated datasets; unspecified parameters remain at default values.

#### 4.1 2D SMLM benchmarking

**DECODE [1].** We used the same parameter settings for model training as in the data simulation (Supplementary Note 2.1).

**SMAP [4].** We used the Difference of Gaussian (DoG) method for peak detection with a scale parameter of $s = 1.2$ and enabled edge exclusion to reduce boundary artifacts. Localization fitting was performed using the Spline method with a pre-calibrated PSF, a 7-pixel ROI, and up to 30 iterations. For low SNR data, $p = 0.45$ was used at densities 3,775 and 7,550, $p = 0.5$ at 18,874, 33,974, and 56,623, and $p = 0.01$ at 113,246. For medium SNR, $p = 0.2$ was used at 3,775, $p = 0.25$ at 7,550, $p = 0.45$ at 18,874 and 33,974, $p = 0.5$ at 56,623, and $p = 0.35$ at 113,246. For high SNR, $p = 0.15$ was used at 3,775, $p = 0.2$ at 7,550, $p = 0.5$ at 18,874, 33,974, and 56,623, and $p = 0.45$ at 113,246.

**CSpline [5].** We used a peak detection radius of 1 pixel with detection thresholds of 0.1 for high and medium SNR and 5 for low SNR. The camera offset was set to 0.1. Fitting used a pre-calibrated PSF. Frame-to-frame peak matching and post-fitting minimum distance filtering were both disabled by setting the matching radius and separation threshold to 0.

**rainSTORM [6].** We used the "Least-Squares Multi Gaussian 2D linear Bg" algorithm without image filtering, initializing the PSF sigma guess at 1.3 pixels. For high SNR, the relative peak threshold was 0.5, sigma range [1, 2] pixels, minimum peak distance 0.1 pixels, and ROI radius 4 pixels. For medium SNR, only the threshold was increased to 2. For low SNR, the threshold was set to 4, sigma range expanded to [0.5, 4], minimum distance increased to 2 pixels, and ROI radius reduced to 3 pixels.

**rapidSTORM [7].** The PSF FWHM was set to 320 nm (X) and 277 nm (Y). Fitting used the Levenberg–Marquardt algorithm with a fixed global threshold. Intensity thresholds of 10, 30, and 50 ADC units were applied for low, medium, and high SNR data, respectively.

#### 4.2 3D SMLM benchmarking

**DECODE [1].** Model training used the same parameter settings as the data simulation (https://srm.epfl.ch/srm/dataset/challenge-3D-simulation/MT0.N2.HD/index.html).

**SMAP [4].** Fitting was performed using a calibrated double-helix PSF with a 25-pixel ROI and $p = 0.06$; all other parameters followed the SMAP configuration used in the 2D SMLM benchmarking.

**CSpline [5].** We used a 10-pixel peak detection radius and a 6.0-sigma threshold. The camera offset was set to 100.0, and a 1-pixel separation threshold was applied post-fitting to filter nearby peaks. Fitting used a bootstrap-calibrated 3D PSF, and peaks were linked across frames using a 0.5-pixel matching radius. The pixel size was set to 100 nm × 100 nm. PSF convolution during peak finding sampled $z$-values at −0.748, −0.374, 0, 0.374, and 0.748 µm.

**QC-STORM [8].** We set the fitting ROI size to 17 pixels and the pixel size to 100 nm × 100 nm. The fitting type was "Double-Helix PSF", with both multi-emitter fitting and low SNR mode enabled. The consecutive fitting radius was set to 80 nm.

**Easy-DHPSF [9].** The conversion gain was set to 1.0 and the pixel size to 100 nm × 100 nm. Processing followed the interactive workflow, where single-PSF images at different axial positions were automatically extracted from SMLM frames and manually screened for clear, symmetric double lobes and good separability from noise. Six frames were selected based on these criteria, shown as frame index and lobe angle: 147 (−50.24°), 115 (−30.08°), 88 (−9.66°), 63 (+10.37°), 38 (+30.50°), and 11 (+49.44°).

#### 4.3 Dynamic 2D SMLM

**DECODE [1].** We configured the camera with a baseline of 100, 0.24 electrons/ADU, 0.81 quantum efficiency, a pixel size of 108 nm × 108 nm, and a read noise sigma of 1.4. For simulation, the average emitter count was set to 46, background intensity was uniformly sampled between 10 and 100, and emitter intensity ranged from 700 to 1750. The `disabled_attributes` parameter was set to 3.

**SMAP [4].** We used $p = 0.5$, with all other settings as in the 2D SMLM benchmarking for SMAP.

**eSRRF [10].** We applied eSRRF with magnification $M = 5$, radius $R = 2$, and sensitivity $S = 2$, using non-overlapping 60-frame temporal segments for reconstruction.

**DBlink [11].** We performed reconstruction with single-molecule localizations from DECODE and SMAP, using the package's pretrained LSTM model configured with a 10-frame trajectory window size, 60-frame non-overlapping processing windows, a pixel size of 108 nm $\times$ 108 nm, and a 4$\times$ super-resolution scale.

#### 4.4 MERFISH analysis

**HSCMERFISH [12].** We used the interactive workflow provided with the official package, using the parameters configured for the evaluated dataset.

#### 4.5 Spike inference benchmarking

**CASCADE [13].** We selected `Global_EXC_40Hz_smoothing50ms_high_noise` from the 155 pretrained models in the official repository, based on testing and parameter matching, to predict spike probabilities. These were then converted into discrete spike trains using the official tool.

**FastL0 [14].** We used the AR(1) model with the calcium decay parameter $\gamma = 1$, consistent with the ground truth used in the simulation. The regularization parameter $\lambda$ was set to 0.005, 0.007, and 0.006 for low-, medium-, and high-SNR data, respectively.

**OASIS [15].** We used the AR(1) model with constrained deconvolution, automatically optimizing the baseline, decay parameter, and minimum spike amplitude from the data. Time points with positive output were converted to spike times using a 0.025 s sampling interval, matching the ground-truth frame rate.

**MLspike [3].** We used the official automatic calibration procedure, guided by ground-truth simulation ranges (Supplementary Note 2.2) for amplitude and time constant, to estimate amplitude, time constant, and noise level. Spike inference was subsequently performed using the calibrated parameters, with the sampling rate set to 40 Hz.

**Peeling [16].** We used ground-truth simulation values (Supplementary Note 2.2) for amplitude, time constant, and noise standard deviation (`noiseSD`). Thresholds were derived from `noiseSD`, with `smtthigh` set to $1.75 \times$ `noiseSD` and `smttlow` set to $-1 \times$ `noiseSD`.

#### 4.6 Background estimation benchmarking

**Dark Sectioning [17].** We used an intensity threshold of 60 to separate foreground from background.

**Sparse Deconvolution [18].** We selected the "Strong background (LI)" mode for background modeling based on empirical testing from the six available modes.

**WBNS [19].** We set the PSF width to 5.5 and the wavelet decomposition level to 0.

**Sliding Paraboloid [20].** The paraboloid radius was set to 0.1 pixels.

**Rolling Ball [21].** The ball radius was set to 10 pixels.

#### 4.7 Active neuron segmentation

We followed the hyperparameter tuning strategy of SUNS [22], selecting parameters from predefined ranges and identifying the optimal combination by maximizing F1 score via four-fold leave-one-out cross-validation on J123's four subvideos and ground truth provided by SUNS.

**SUNS batch [22].** We searched over SNR threshold $[48.7, 97.3]$, minimum area $[0.2, 0.51]$ $\mu$m$^2$, probability threshold $[3.1, 6.2]$, center-of-mass distance threshold $[1, 7]$ $\mu$m, and minimum consecutive frames $[1, 7]$.

**STNeuroNet [23].** We used a training block size of $144 \times 144 \times 256$ with 25,000 iterations. The minimum area was tuned over $[50, 140]$ pixels and the probability map threshold over $[0.9, 1.0]$.

**CaImAn batch [24].** We set the calcium transient duration to 0.5 s and the number of components per patch to 20. Tuned parameters included spatial correlation threshold $[0.7, 0.95]$, minimum SNR $[3, 6]$, CNN upper threshold $[0.8, 0.95]$, and lower threshold $[0, 0.6]$.

**Suite2p [25].** We set the calcium transient time constant to 0.5 s. The ROI detection threshold scaling factor was searched over $[0.6, 2.0]$.

### 5 Supplementary tables

**Table 1**: Parameters for test data simulation using DECODE

| **Category and parameter** | **Value** |
|---|---|
| **Camera** | |
| Baseline | 398.6 |
| Convert to Photons | true |
| Electrons per ADU | 5.0 |
| EM Gain | 100 |
| Pixel Size | (127.0, 117.0) |
| Quantum Efficiency (QE) | 1.0 |
| Read Noise Sigma | 58.8 |
| Spurious Noise | 0.0015 |
| **Simulation** | |
| Background Uniform | (20.0, 200.0) |
| Density | *manually set* |
| Emitter Average | *manually set* |
| Emitter Extent | $[[-0.5, 511.5], [-0.5, 511.5], [-1, 1]]$ |
| Image Size | (512, 512) |
| Intensity (Mean, Sigma) | *manually set* |
| Intensity Threshold | null |
| Lifetime Average | 1.0 |
| Mode | acquisition |
| Photon Range | null |
| PSF Extent | $[[-0.5, 511.5], [-0.5, 511.5], \text{null}]$ |
| ROI Auto Center | false |
| ROI Size | null |
| XY Unit | px |

**Table 2**: Parameters for test data simulation using MLspike

| **Category and parameter** | **Value** |
|---|---|
| **Spike train generation** | |
| Spike rate | *manually set* |
| Mode | bursty |
| Spikes per burst (mean) | 1 |
| Inter-spike interval within burst (mean) | 0.01 s |
| **Acquisition** | |
| Sampling interval ($dt$) | 0.025 s (40 Hz) |
| Total duration ($T$) | 30 s |
| **Calcium dynamics** | |
| Model type | 1exp |
| Amplitude per spike ($a$) | *manually set* |
| Decay time constant ($\tau$) | *manually set* |
| Delay | 0 |
| Saturation level | 0.1 |
| Hill coefficient | 1 |
| Calcium baseline ($c_0$) | 0 |
| Fluorescence baseline ($F_0$) | 1 |
| Initial calcium state ($x_0$) | 0 |
| **Noise and drift** | |
| Noise level ($\sigma$) | *manually set* |
| Drift method | Basis functions |
| Drift effect | Multiplicative |
| Drift harmonics | 3 |
| Drift amplitude | 0.005 |

#### 6 Supplementary figures

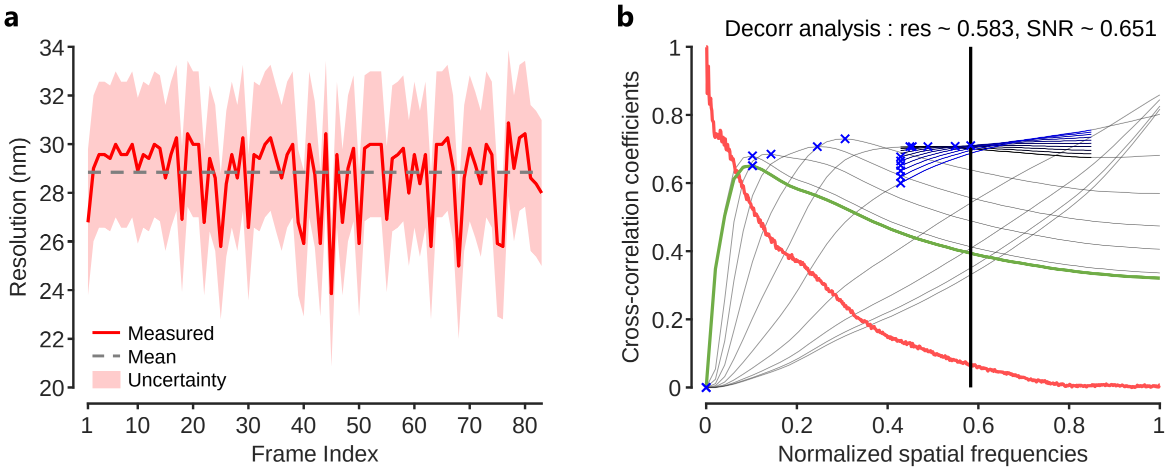

**Supplementary Fig. 1. Decorrelation analysis for spatial resolution quantification. a,** The results of the decorrelation analysis algorithm [26] for each frame of the super-resolution video rendered with DEPAF (Supplementary Video 1). Red line, measured spatial resolution of each frame; gray dashed line, mean spatial resolution across all frames; red shaded region, the $\pm 3$ nm uncertainty range reported in Ref. [26]. **b,** Output of the decorrelation analysis algorithm for the frame with the lowest spatial resolution (Frame 77). The results show that the maximal spatial frequency in our reconstruction $k_c$ corresponds to $\sim 58\%$ of the maximal achievable frequency in our system. According to the spatial resolution estimation formula provided in Ref. [26]: $d = \frac{2p}{k_c}$, where $d$ is the spatial resolution and $p$ is the pixel size (9 nm in this case), the spatial resolution of this frame is $\sim 31$ nm. Green line, decorrelation functions before high-pass filtering; red line, radial average of the log of the absolute value of the Fourier transform of the analyzed frame; gray lines, all high-pass filtered decorrelation functions; blue to black lines, decorrelation functions with refined mask radius and high-pass filtering range; blue crosses, all local maxima; black vertical line, cutoff frequency $k_c$.

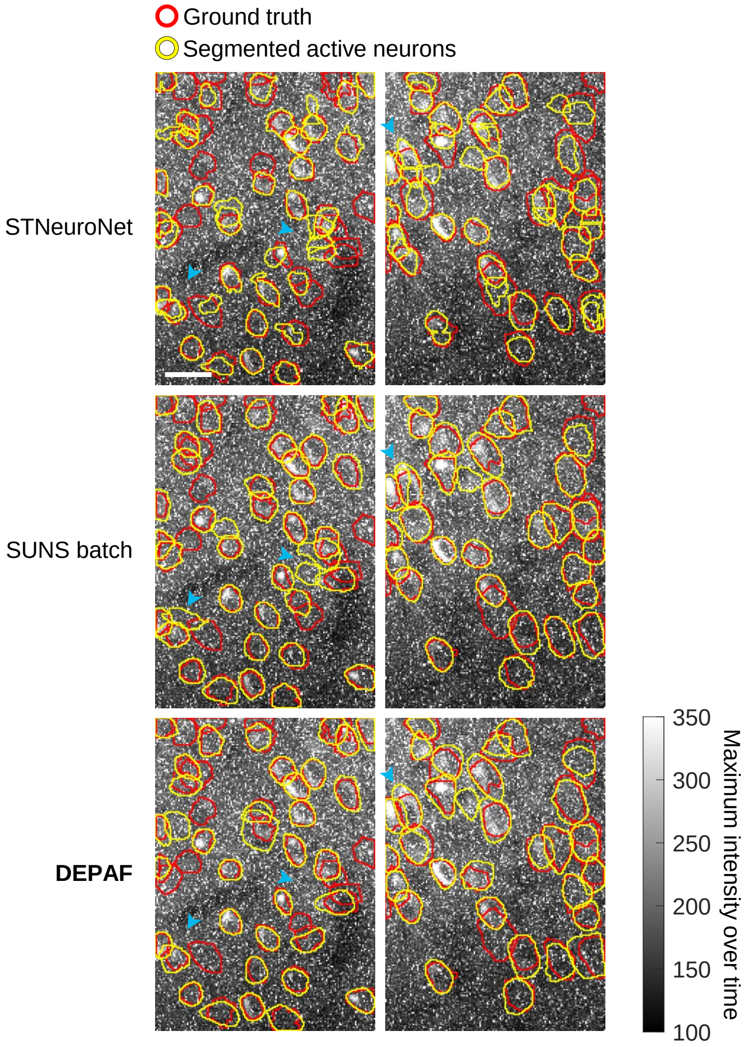

**Supplementary Fig. 2. Comparison of DEPAF and *supervised* methods for active neuron segmentation.** Example segmentation results of DEPAF, STNeuroNet [23], and SUNS batch [22] from the second and fourth quadrants of video J123, overlaid on the time-axis maximum intensity projection. Blue arrows indicate regions of difference. Scale bar, 20 $\mu$m.

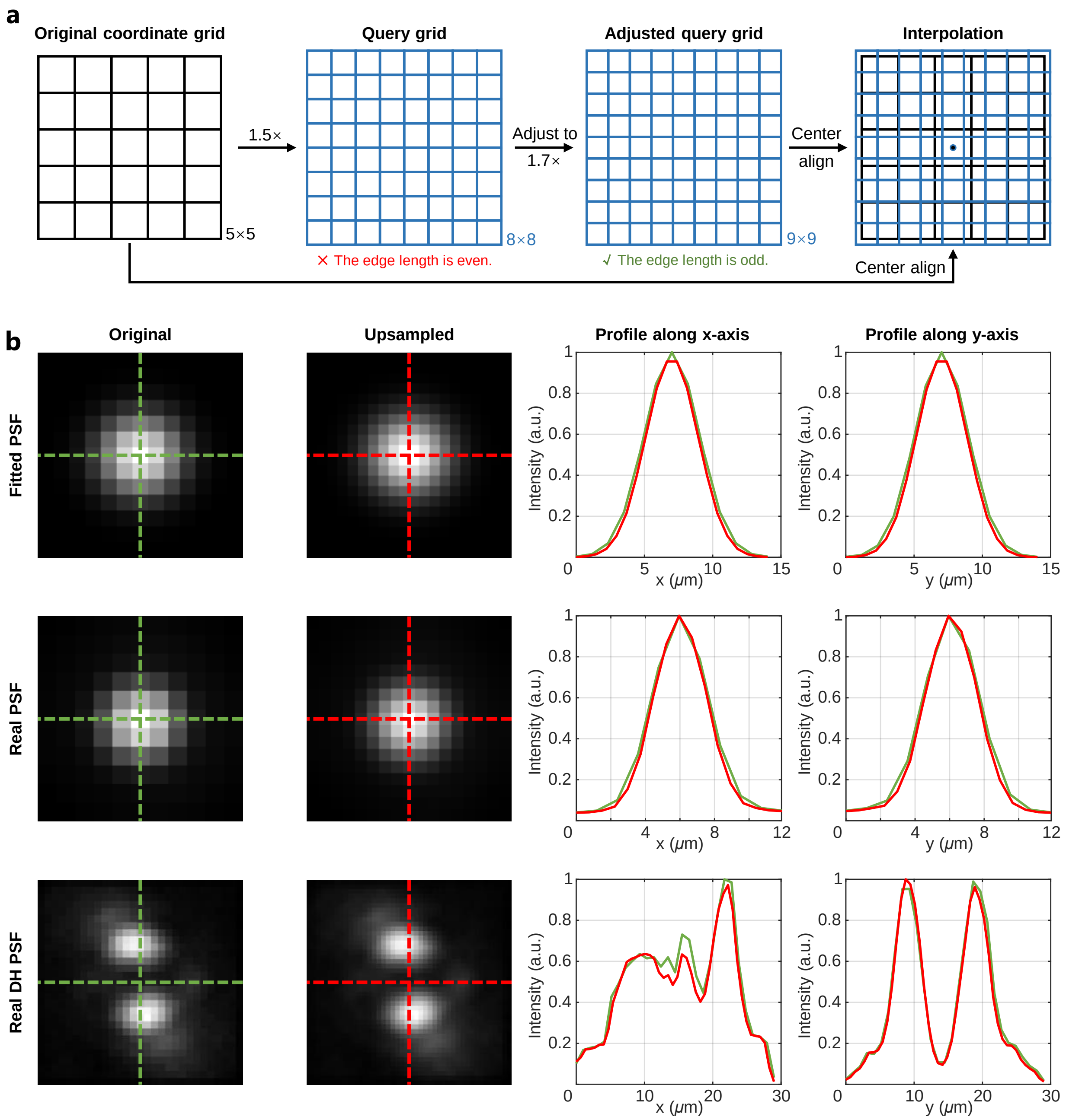

**Supplementary Fig. 3. Schematic depiction of the center-aligned upsampling technique and its application results. a,** A query grid is generated from the original coordinate grid and adjusted to have an odd edge length. Center alignment is achieved through coordinate mean matching, followed by interpolation for upsampling. **b,** Results for fitted PSF (first row), real PSF (second row), and real double-helix (DH) PSF (third row). The first column shows the original data, the second column shows the upsampled data, and the third and fourth columns compare intensity profiles along the $x$-axis and $y$-axis (dashed-line positions), with green representing the original and red the upsampled data.

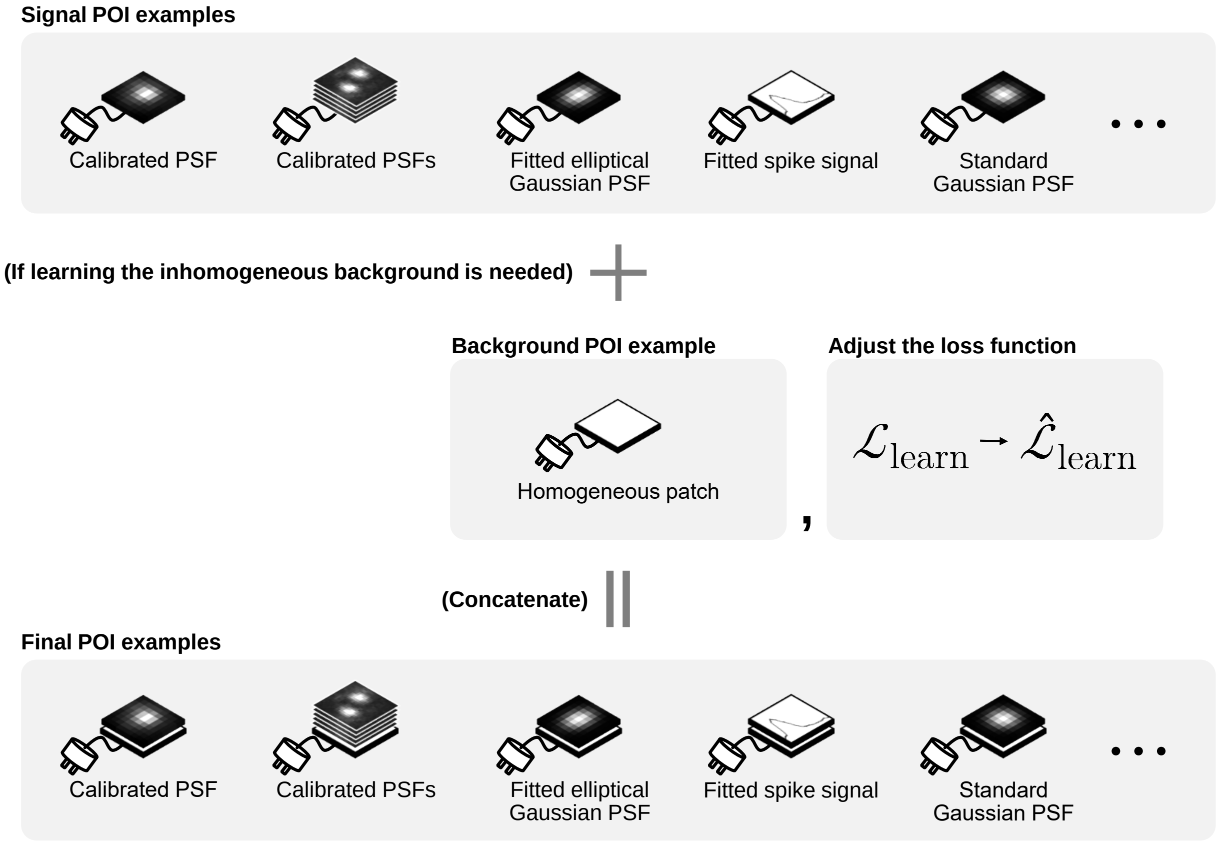

**Supplementary Fig. 4. Schematic depiction of the inhomogeneous background learning technique.** If learning the inhomogeneous background is needed, DEPAF automatically adds a background learning term to the loss function and introduces an additional homogeneous patch as a background POI example alongside the original signal POI examples. Subsequently, by concatenating the signal POI examples with the background POI example along the channel dimension, the final POI examples used by the DEPAF model is generated. Through this mechanism, the DEPAF model can adaptively learn and capture arbitrary inhomogeneous backgrounds present in the data.

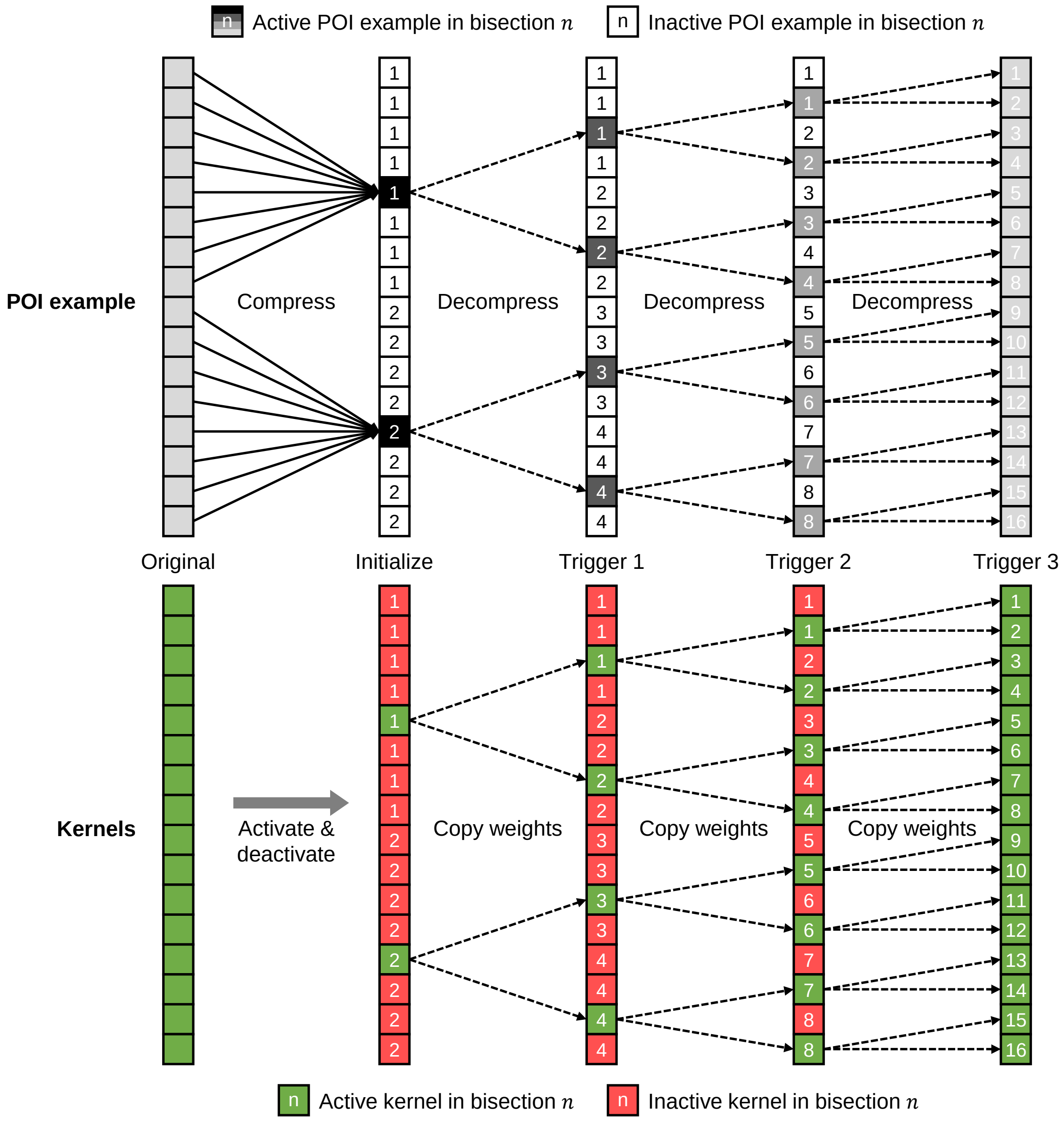

**Supplementary Fig. 5. Schematic depiction of the POI diffusion technique.** The technique divides the POIs in the POI example into two groups via bisection, compressing each group into a central channel using mean pooling. In $\mathcal{M}_{\text{net}}$'s final convolutional layer, only kernels corresponding to the central channels are activated for training. When validation loss plateaus, POI diffusion triggers: recursively applying bisection to determine new activated channels and decompressing POIs to nearest active channels. Newly activated kernels inherit weights from previously activated kernels via nearest copy and join training. This process iterates until all channels are activated.

#### 7 Supplementary video descriptions

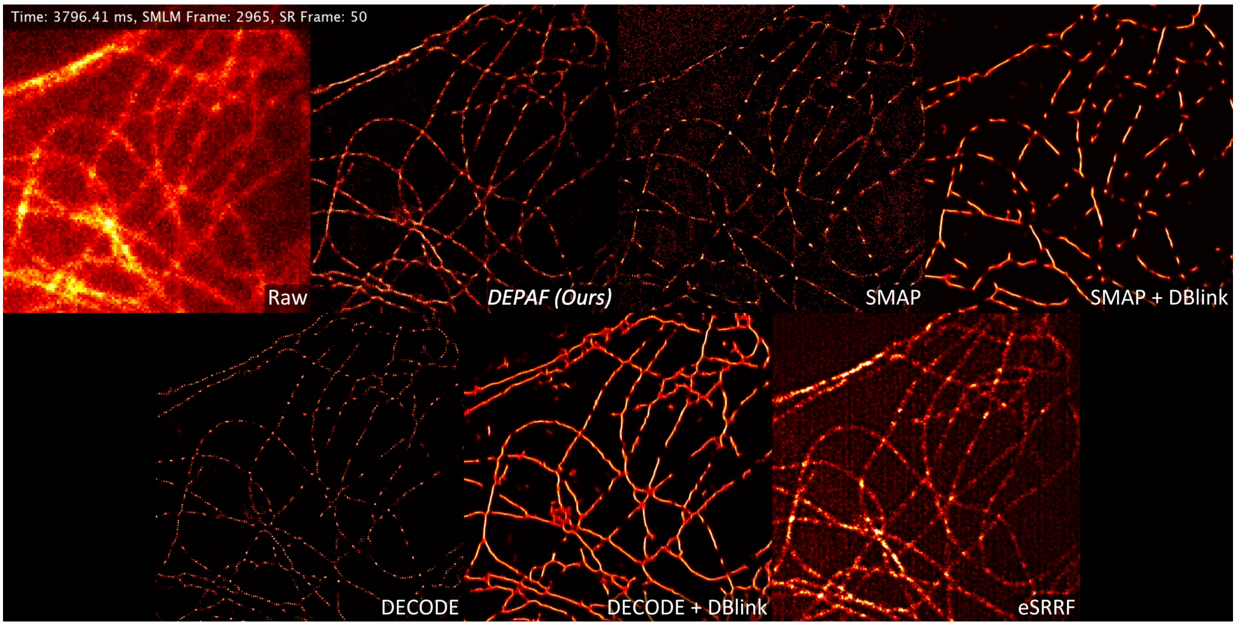

**Supplementary Video 1. Method Comparison for Super-Resolution Imaging of Cytoskeletal Fluctuations.** This video demonstrates the visualization of cytoskeletal fluctuations using the DEPAF method presented in this work, along with five other comparative methods. The SMAP method [4], under strong interference, struggles with detecting and localizing high-density PSFs, leading to low spatiotemporal resolution with high noise and missing details. When combined with DBlink [11], the sparse localizations from SMAP cause the spatiotemporal interpolation to introduce artifacts and missing details. Although the DECODE method [1] can handle high-density PSF detection, it struggles to localize weak PSFs in ultra-high-density scenarios, causing grid artifacts and loss of detail. Combining DECODE with DBlink improves temporal resolution but introduces artifacts and noise due to spatiotemporal interpolation based on the suboptimal results of DECODE. The eSRRF method [10], being a non-SMLM technique, is suitable for fast imaging but suffers from lower spatial resolution ($\sim$70 nm) and high sensitivity to noise, limiting its ability to accurately reconstruct rapidly changing subcellular structures. In contrast, the DEPAF method, proposed in this work, utilizes ultra-high-density blinking dSTORM imaging with a high-speed camera ($\sim$781 FPS), achieving 76.8 ms temporal resolution ($\sim$13 FPS) and $\sim$30 nm spatial resolution, enabling precise capture of rapid, subtle cytoskeletal dynamics with a balanced spatiotemporal resolution.

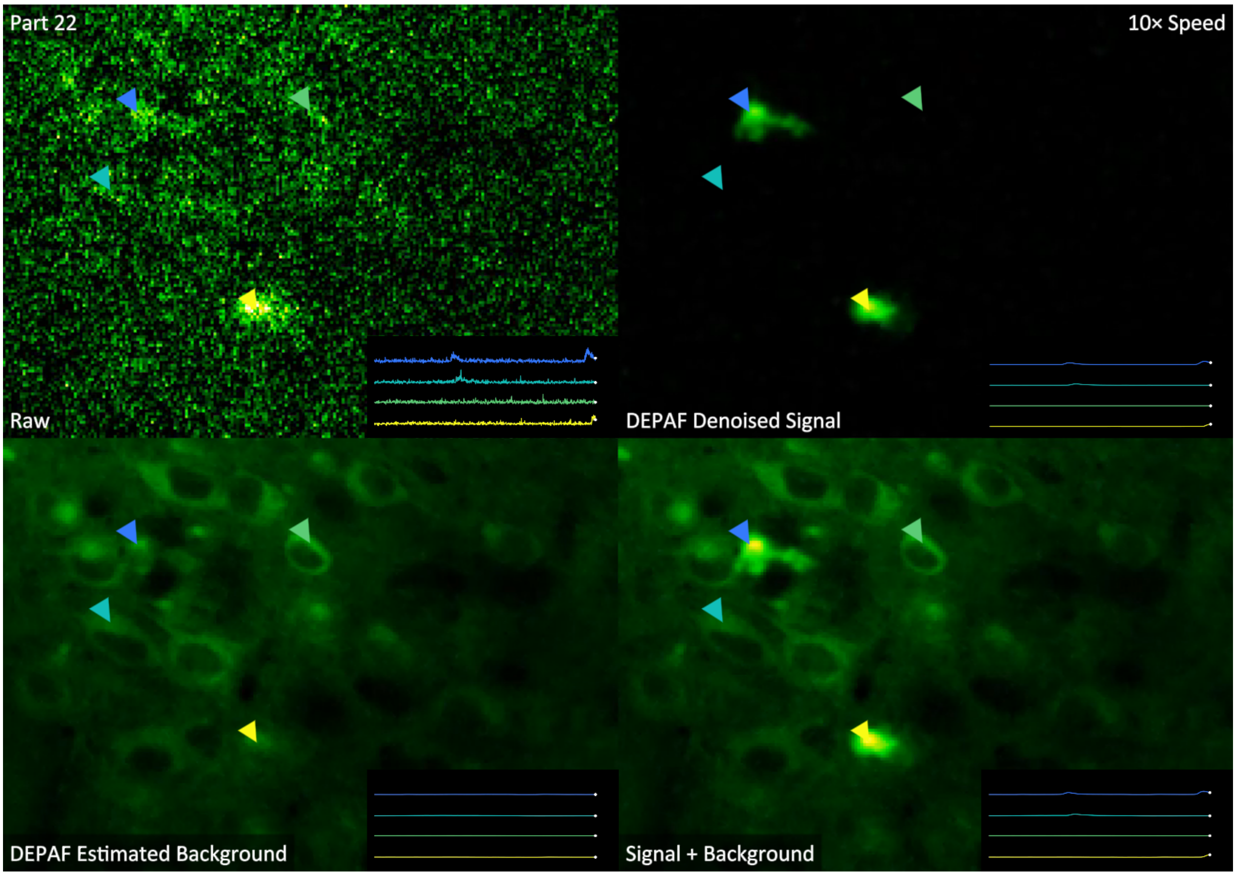

**Supplementary Video 2. Denoising with Background Separation Using DEPAF in J123 Video from the CalmAn Dataset (Preview).** This video demonstrates denoising and background separation of the J123 video, a representative low-SNR two-photon calcium imaging video from the CaImAn dataset [24], using the DEPAF method. DEPAF treats each pixel's temporal signal as a one-dimensional (1D) image and uses a fitted spike signal to accurately estimate the positions and amplitudes of overlapping, weak spikes while effectively separating spatiotemporally heterogeneous backgrounds. The result is a high-fidelity denoised video that retains only active neuronal signals. This video is a preview, the full-length version is available at https://zenodo.org/records/14145974.
